# Next Generation TB Drug Combinations from the Pan-TB Consortium: Combination Efficacy and Contributions of Individual Agents, Evaluated in a BALB/c Mouse TB Model

**DOI:** 10.64898/2026.06.18.733138

**Authors:** Sylvie Sordello, Alain Le Coupanec, Zoï Vahlas, Chiara Roversi, Roberto Visentin, Xavier Boulenc, Denise Federico, Simone Zannoni, Simone Modolo, Augusto Celon, Roberto Petterlini, Cécile Pascal, Fanny Deglave, Alessia Tagliavini, Marco Pergher, Khisimuzi Mdluli, Micha Levi, Todd A. Black, Robert H. Bates, Yongge Liu, Yohei Hayashi, Clara Aguilar Perez, Hermann David, Debra Flood, Anna M. Upton

**Affiliations:** Translational Biology, Infection Diseases, Evotec France, Toulouse, France; Pharmacometrics, Aptuit (an Evotec Company), Verona, Italy; DMPK, Evotec Infectious Diseases Lyon, Lyon, France; DMPK, Evotec France, Toulouse, France; Gates Medical Research Institute, Massachusetts, USA; TB Alliance, New York, USA; GSK Global Health Medicines R&D, Tres Cantos, Spain; Otsuka Pharmaceutical Development and Commercialization, Inc., Rockville, USA; Otsuka Pharmaceutical Co., Ltd., Tokyo, Japan; Johnson & Johnson, Beerse, Belgium; Gates Foundation, Seattle, Washington, USA; Evotec USA inc., New Jersey, USA

## Abstract

The Project to Accelerate New Treatments for Tuberculosis (PAN-TB) aims to accelerate development of shorter, simpler and safer pan-TB combinations. We previously identified 3 out of 25 first-generation novel PAN-TB 4-drug combinations, that cured 90% of mice in less than 3 months, at clinically relevant doses in the relapsing mouse model of TB. These regimens include BPa830Sut, BPa286Sut and BQSut286 (B: bedaquiline; Pa: pretomanid; 830: GSK3211830; 286: GSK2556286; Sut: sutezolid; Q: quabodepistat). Here, we assess the efficacy of these combinations where the original candidates are substituted next-generation or more advanced compounds (ganfeborole (656) for 830, sorfequiline, S for B, TBD09 for Sut and TBD11 for 286) and the individual contributions of specific agents. Six novel regimens demonstrated bactericidal activity more rapid than comparators PHMZ (Rifapentine P, Isoniazid H, Moxifloxacin M, Pyrazinamide Z) and BPaMZ. Modelled cure/relapse data showed that SPa286Sut, SPaSut and SPa656Sut cured 90% of mice in about 1 month, while SPa286, SPaQTBD11 and SPaTBD09 in less than 2 months, faster than PHMZ. Consistent with our previous findings, the fastest-curing regimens centered on a diarylquinoline (S), a nitroimidazole (Pa) and an oxazolidinone (TBD09 or Sut) together with an Rv1625c agonist (TBD11 or 286), DprE1 inhibitor (Q) or a LeuRS inhibitor (656). Notably, significant contributions to sterilizing efficacy were demonstrated for S in all combinations and for Pa, Sut, TBD09, Q and TBD11 or 286 in specific S-containing combinations. These findings suggest potential for these novel agents and combinations to improve treatment of both DS-and DR-TB.

## INTRODUCTION

Tuberculosis (TB) currently ranks as the number one infectious agent killer [1]. Of the estimated 10.7 million new cases reported in 2024, 390,000 involved rifampicin resistant or multidrug resistant (RR/MDR) TB. Although treatments are available for both drug-susceptible (DS) and RR/MDR TB, they are long, complex, and poorly tolerated. Additionally, rapid drug susceptibility testing (DST) is required to ensure appropriate treatment. Access to both DST and RR/MDR TB treatment options are currently limited, presenting barriers to TB control and compromising patient outcomes [2].

Novel TB drug combinations aligned to the World Health Organization (WHO) pan-TB target regimen profile would eliminate the need for DST. These drug combinations are defined as first-line regimens that could be used without prior knowledge of a TB patient’s drug-resistance profile [3]. Introducing shorter, safer and simpler pan-TB regimens to high TB burden countries could result in substantial benefits to patients and health systems [4]. Although progress has been made towards treatments that fit WHO target regimen profiles for drug-susceptible TB (DS-TB) and rifampicin resistant TB (RR-TB) [3], more limited progress has been made towards development of pan-TB regimens.

The Project to Accelerate New Treatments for TB (PAN-TB), a philanthropic, non-profit and private sector collaboration (<u>Tuberculosis Prevention | PAN-TB</u>), was formed with the aim to accelerate prioritization and development of promising pan-TB combinations. The Collaboration conducts nonclinical and clinical studies utilizing the assets and technologies of its members.

We previously reported the pharmacokinetics (PK) and comparative efficacy of 25 unique PAN-TB 4-drug combinations, evaluated in a BALB/c relapsing mouse model (RMM) of TB. These combinations were comprised of 8 agents with pan-TB potential: bedaquiline (B), an ATP synthase inhibitor [5], delamanid (Del) and pretomanid (Pa), nitroimidazole mycolic acid synthesis inhibitors [6–7], quabodepistat (Q) an inhibitor of decaprenylphosphoryl-β-D-ribose 2’-epimerase [DprE1] [8], GSK3211830 (830) and ganfeborole (GSK3036656, 656), two oxaborole leucyl-tRNA synthetase (LeuRS) inhibitors [9], GSK2556286 (286) an Rv1625c agonist, c-AMP mediated cholesterol catabolism inhibitor [10] and sutezolid (Sut) an RNA translation inhibitor [11].

Of the 25 PAN-TB combinations tested, 15 cured 90% of mice more rapidly than the standard of care for DS-TB, RHZE/RH (Rifampicin, Isoniazid, Pyrazinamide, Ethambutol) and the best-performing PAN-TB combinations (BPa830Sut, BPa286Sut and BQSut286) cured 90% of mice in less than 3 months [12]. These 3 top-ranked 4-drug combinations are centered on a diarylquinoline (B)/oxazolidinone (Sut) core. Next generation diarylquinolines and oxazolidinones have now entered development and demonstrate improved potency and/or safety and PK profiles. Accordingly, the PAN-TB consortium has prioritized evaluation of the next generation diarylquinoline sorfequiline (S) (formerly TBAJ-876) [13] as well as the oxazolidinone TBD09 (formerly MK-7762) [14]. In addition, TBD11 (formerly mCLB-073), an activator of adenylyl cyclase encoded by Rv1625c [15, 16] has been added to PAN-TB’s priority agents, as a replacement for 286 which was discontinued during Phase I clinical evaluation. Previous studies conducted by others in the BALB/c RMM, have demonstrated treatment-shortening potential for these novel agents within specific drug combinations [14, 17]

Here, using a similar BALB/c RMM workflow to that presented in our previous work, we aimed to evaluate the impact of substituting S for B in the PAN-TB combinations previously identified as being most effective and we sought to determine the relative contributions to bactericidal and sterilizing efficacy of compounds within shared chemical classes or mechanisms of action including TBD09 and TBD11 in place of the previously-studied Sut and 286. Finally, we aimed to quantify the treatment-shortening effects of S, TBD11, 286, Sut, TBD09, Q, and Pa within the studied combinations. In these studies, we compared time to cure of the novel PAN-TB combinations to that demonstrated in the BALB/c RMM by the clinical benchmark PHMZ (where P is Rifapentine) which cured DS-TB patients more rapidly than RHZE/RH [18]. We also compared the efficacy of the novel regimens to that of the experimental regimen BPaMZ (SimpliciTB) which has demonstrated treatment-shortening potential in this mouse TB model, as well as in patients relative to RHZE/RH [19].

## RESULTS

### Six PAN-TB drug combinations demonstrate bactericidal activity in mice that is more rapid than BPaMZ

All PAN-TB combinations tested **(Table 1)** showed significant time-dependent bactericidal activity, over the course of 8 weeks’ treatment, reducing lung CFU to undetectable levels, with the exception of SQ286TBD09 where colonies were still detectable in one mouse (**Figure 1A and 1B**). Similar bactericidal kinetics were observed between equivalent combinations that contained B compared with S and when 830 was replaced by 656, as well when Pa was replaced by Del (**Figure 1A**).

**Figure 1.**
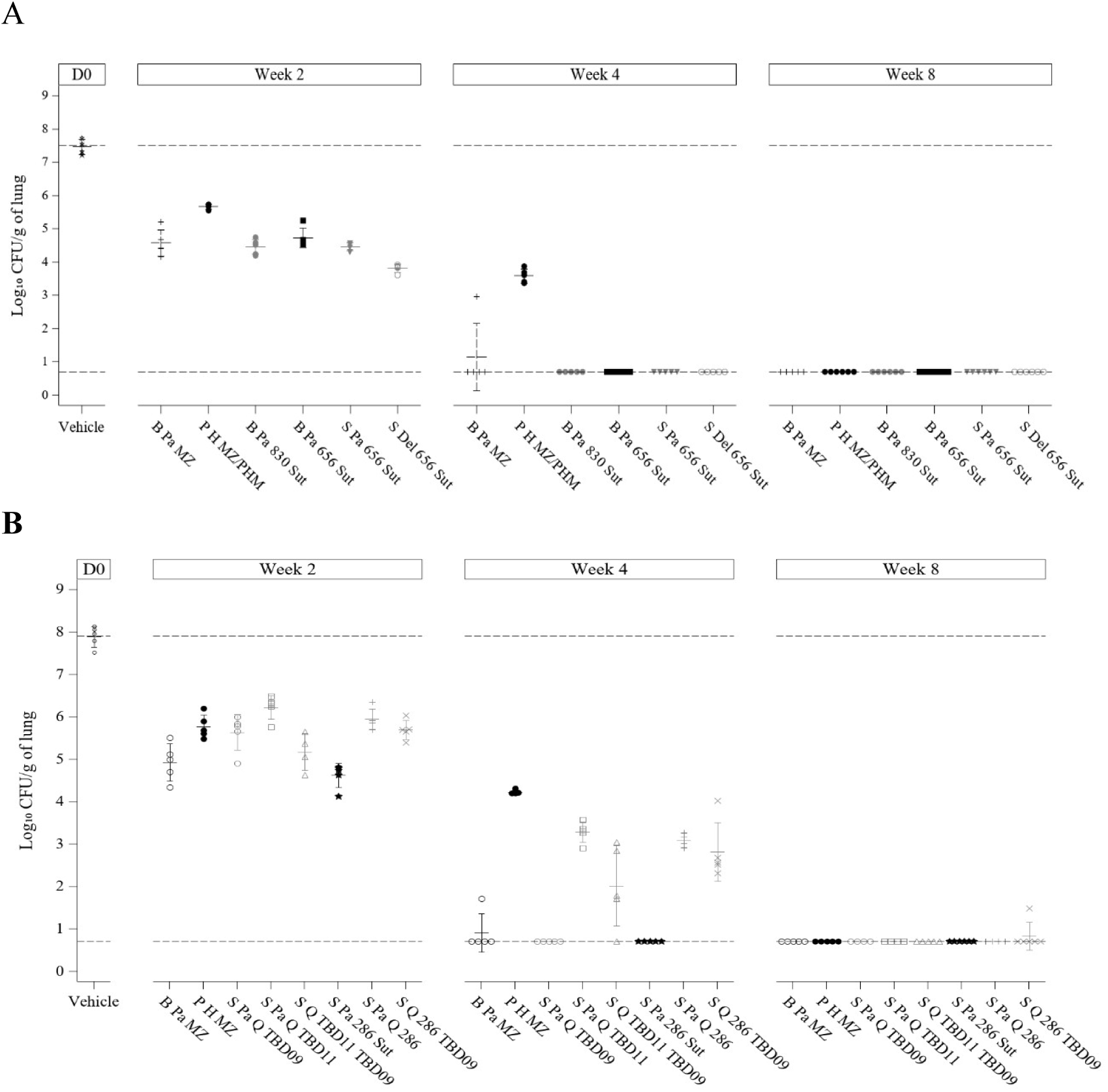
Lung bacterial burden at the end of treatment in Study 1 (A) and Study 2 (B) studies. Lung CFU/ g of lung, of BALB/c mice intranasally infected with *M.tb* H37Rv, after different durations of oral treatment dosed 5/7. Treatments were initiated 2-weeks post infection. (mean Log_10_ CFU/ g of lung +/- sd).

**Table 1.** PAN-TB selected 3- and 4-drug combinations. Combinations were tested in Study 1 and Study 2 of the PAN-TB collaboration.

| **Study 1** | **Study 2** |
| --- | --- |
| B Pa M Z | S Pa 286 Sut |
| S Pa 656 Sut | B Pa M Z |
| B Pa 656 Sut | *S Pa Sut* |
| S Del 656 Sut | *S Pa 286* |
| B Pa 830 Sut | S Pa Q TBD11 |
| P H M Z | *S Pa TBD09* |
|  | S Q TBD11 TBD09 |
|  | S Pa Q TBD09 |
|  | S Pa Q 286 |
|  | S Q 286 TBD09 |
|  | *S Pa TBD11* |
|  | *S Pa Q* |
|  | P H M Z |
|  | *S Q TBD09* |
| 286, GSK2556286; 656, GSK3036656; 830, GSK3211830; B, Bedaquiline; Del, Delamanid; H, isoniazid; M, moxifloxacin; Pa, Pretomanid; P, Rifapentine; Q, Quabodepistat; Sut, Sutezolid; Z, pyrazinamide; S Sorfequiline |  |

Six 4-drug combinations (BPa830Sut, BPa656Sut, SPa656Sut, SPa286Sut, SDel656Sut and SPaQTBD09), reduced lung CFU to undetectable levels after 4 weeks’ treatment (**Figure 1A and B**), more rapidly than either BPaMZ (1/5 mice remained positive after 4 weeks’ treatment) or PHMZ, which reduced lung CFU by only 4 Log_10_ CFU (5/5 mice positive) in this timeframe. Both BPaMZ and PHMZ required 8 weeks’ treatment to render mice culture-negative.

Comparisons of 3 and 4-drug combinations were performed, to evaluate contributions of individual agents – i.e. as the result of adding a 4^th^ drug to a 3-drug combination. Sut, TBD09, Pa and TBD11 contributed significant bactericidal activity within tested 4-drug combinations: Addition of Sut to SPa286; TBD09 to SPaQ; Pa to SQTBD09, and TBD11 to SQTBD09 resulted in additional reductions in lung CFU of 2.5, 2.6, 3 or 1.7 Log_10_ after 4 weeks of treatment respectively (**Table S2 A and B**, **Table 2**). Substituting TBD11 for 286 in SQTBD09, SPaQX and SPaX, did not result in a change to the bacterial burden after 4 weeks’ treatment (**Table S2 A and B**, **Table 3**).

**Table 2.**
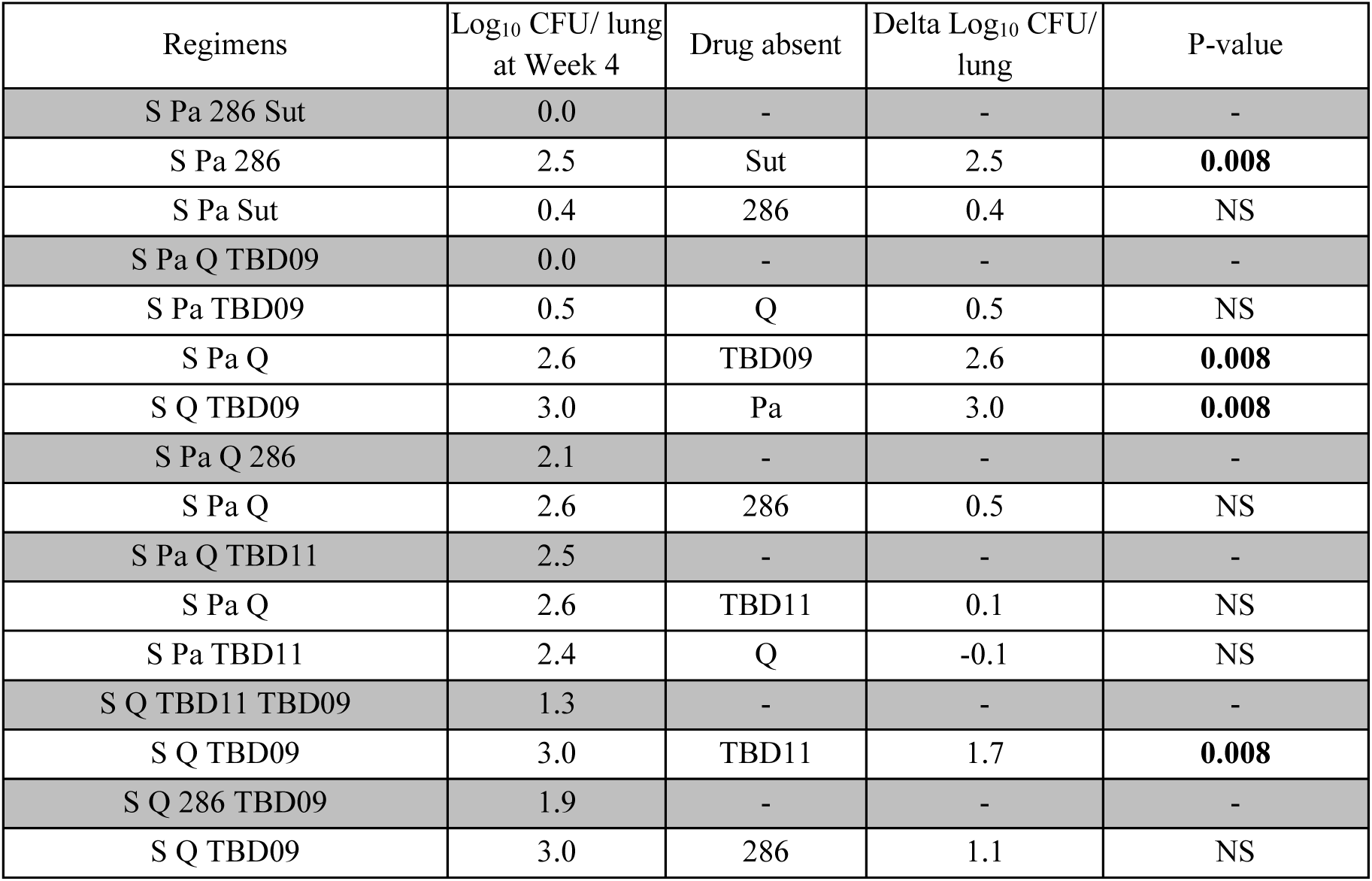
Contribution of drugs to Bactericidal activity after 4 weeks’ treatment: Comparison of 3- and 4-drug combinations.

**Table 3.**
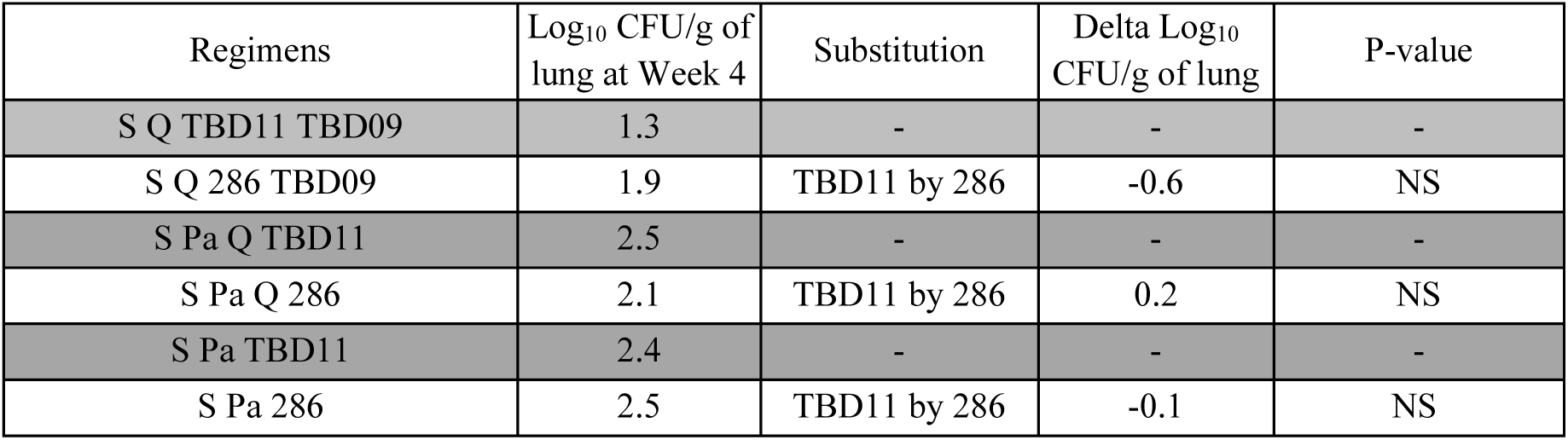
Comparison of 286 and TBD11 contributions to bactericidal activity after 4 weeks’ treatment.

### Five next generation combinations cured 90% of mice in less than 2 months, with sorfequiline driving the regimen performance

Quantifying the proportion of mice that were culture positive (i.e. exhibiting relapses) twelve weeks after the end of treatment (**Table 4)** showed that BPaMZ and SPa656Sut sterilized mice with 6 weeks’ treatment (**Table 4A**), whereas SPa286Sut out-performed BPaMZ, displaying 0% relapse after 4 weeks’ treatment (**Table 4B**). With the exception of SPaQTBD11, which sterilized mice with 8 weeks treatment, all other tested PAN-TB 4-drug combinations (BPa656Sut, SDel656Sut, BPa830Sut, SQTBD11TBD09, SPaQTBD09, SPaQ286 and SQ286TBD09) sterilized mice with 10 weeks’ treatment, at least as rapidly as PHMZ/PHM (**Table 4A and 4B**).

**Table 4.**
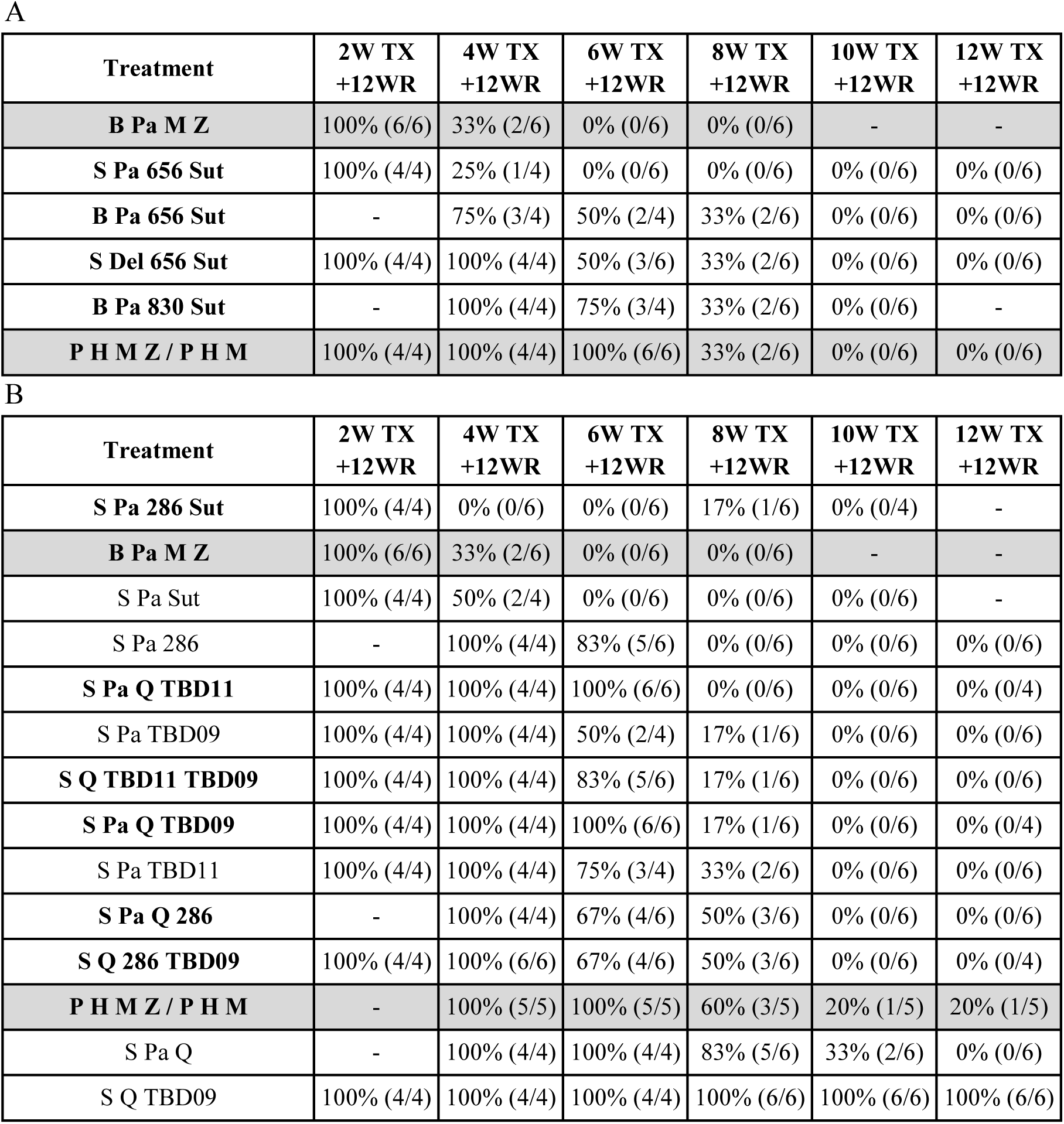
Relapse percentage for study Study 1 (A) and Study 2 (B) ranked based on efficacy % of relapse (positive culture mice/total number of mice), Treatment (TX), Weeks (W), Relapse (R).

To predict the treatment duration needed for cure, we applied a logistic Emax model developed a population approach that has demonstrated good performance in predicting the relapse profiles of the 4-drug combinations, especially for comparator BPaMZ for which data from different studies and labs are available (**Figure S1**). Individual relapse-time curves for each combination are shown in **Figure 2** and the corresponding estimated population time to 50% cure (T50) and derived time to 90% cure (T90) values are shown in **Table 5**.

**Figure 2.**
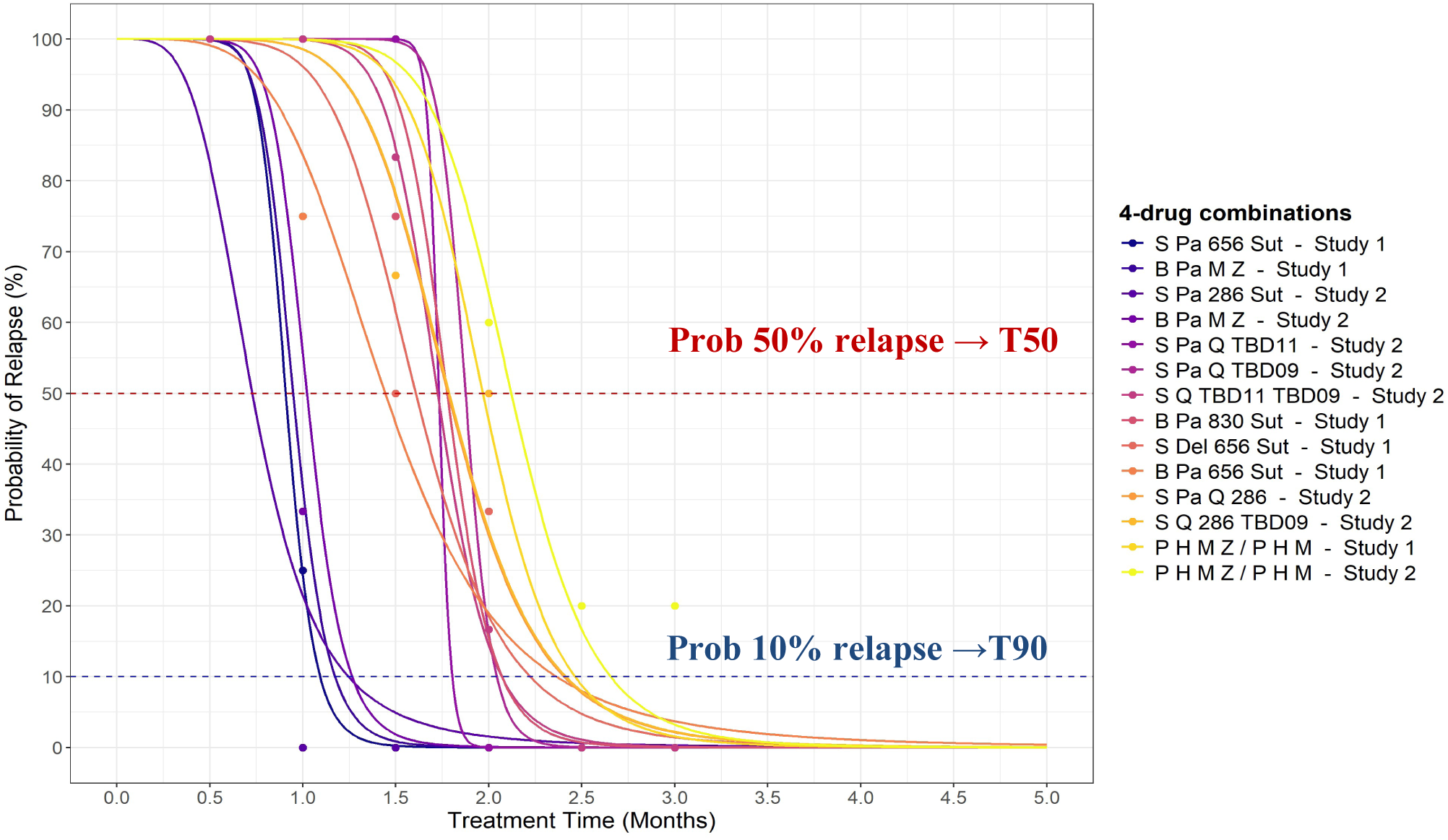
Probability of relapse with treatment duration for BALB/c mice treated with 4-drug combinations tested in Study 1 and Study 2. Sterilization curves indicating the probability of relapse over treatment time, derived by fitting observed relapse data (Table 3) with an Emax model developed using a large historical RMM dataset. Observed relapse data are indicated for each test regimen using dots. The time to 50% cure/relapse (T50) is estimated from these curves, and the time to 90% cure (T90) is derived utilizing time to 50% cure estimates together with steepness of the curve (gamma parameter) as explained in Methods.

**Table 5.** Estimated population T50 and derived population T90 (with 95% confidence interval) for 4-drug regimens tested in Study 1 and Study 2.

| Regimen | Estimated<br>Population T50<br>(Months) | Derived<br>Population T90<br>(Months) [95% CI] |
| --- | --- | --- |
| **S Pa 286 Sut** | 0.48 | 0.82 [0.69, 0.96] |
| S Pa Sut | 0.83 | 0.96 [0.84, 1.07] |
| **S Pa 656 Sut** | 0.80 | 0.97 [0.89, 1.04] |
| **B Pa M Z** | **0.86** | **1.06 [0.99, 1.14]** |
| S Pa 286 | 1.60 | 1.74 [1.58, 1.91] |
| **S Pa Q TBD11** | 1.77 | 1.84 [1.67, 2.01] |
| S Pa TBD09 | 1.54 | 2.00 [1.81, 2.20] |
| **S Q TBD11 TBD09** | 1.76 | 2.11 [1.91, 2.31] |
| **S Pa Q TBD09** | 1.95 | 2.12 [1.93, 2.31] |
| S Pa TBD11 | 1.81 | 2.30 [2.09, 2.51] |
| **B Pa 830 Sut** | 2.00 | 2.33 [2.12, 2.54] |
| **S Del 656 Sut** | 1.77 | 2.44 [2.30, 2.58] |
| **S Pa Q 286** | 1.83 | 2.47 [2.24, 2.70] |
| **S Q 286 TBD09** | 1.84 | 2.48 [2.25, 2.71] |
| **B Pa 656 Sut** | 1.54 | 2.52 [2.37, 2.67] |
| **P H M Z/P H M** | **2.26** | **2.83 [2.35, 3.32]** |
| S Pa Q | 2.51 | 2.98 [2.72, 3.24] |
| S Q TBD09 | NC | NC |

T90 values for the 10 analyzed novel combinations ranged from 0.8 to 2.8 months where the T90 for the PHMZ/PHM comparator was the highest (2.8 months) (**Table 5** and **Figure 2**). SPa286Sut (T90 of 0.82 months) out-performed BPaMZ (T90 of 1.06 months) by 0.24 months (p = 0.002), whereas SPa656Sut (T90 of 0.97 months) performed similarly to BPaMZ (p=0.079). All three of these regimens cured 90% of mice in approximately 1 month, while SPaQTBD11 exhibited a T90 of less than 2 months (1.84 months) (**Table 5**). Two additional 4-drug combinations, SQTBD11TBD09 (2.1 months) and SPaQTBD09 (2.1 months) cured 90% of mice in approximately 2 months, as rapidly as BPaL (2.3 months) (**Figure 3**). All other tested combinations, namely BPa830Sut, SDel656Sut, SPaQ286, SQ286TBD09 and BPa656Sut cured 90% of mice in less than 2.5 months (**Table 5**). Notably, 3-drug regimens such as SPaSut, SPa286 and SPaTBD09 cured mice less than 2 months, as rapid as some 4-drug regimens such as BPaMZ, SPaQTBD11 and SQTBD11TBD09. (**Table 5**).

**Figure 3.**
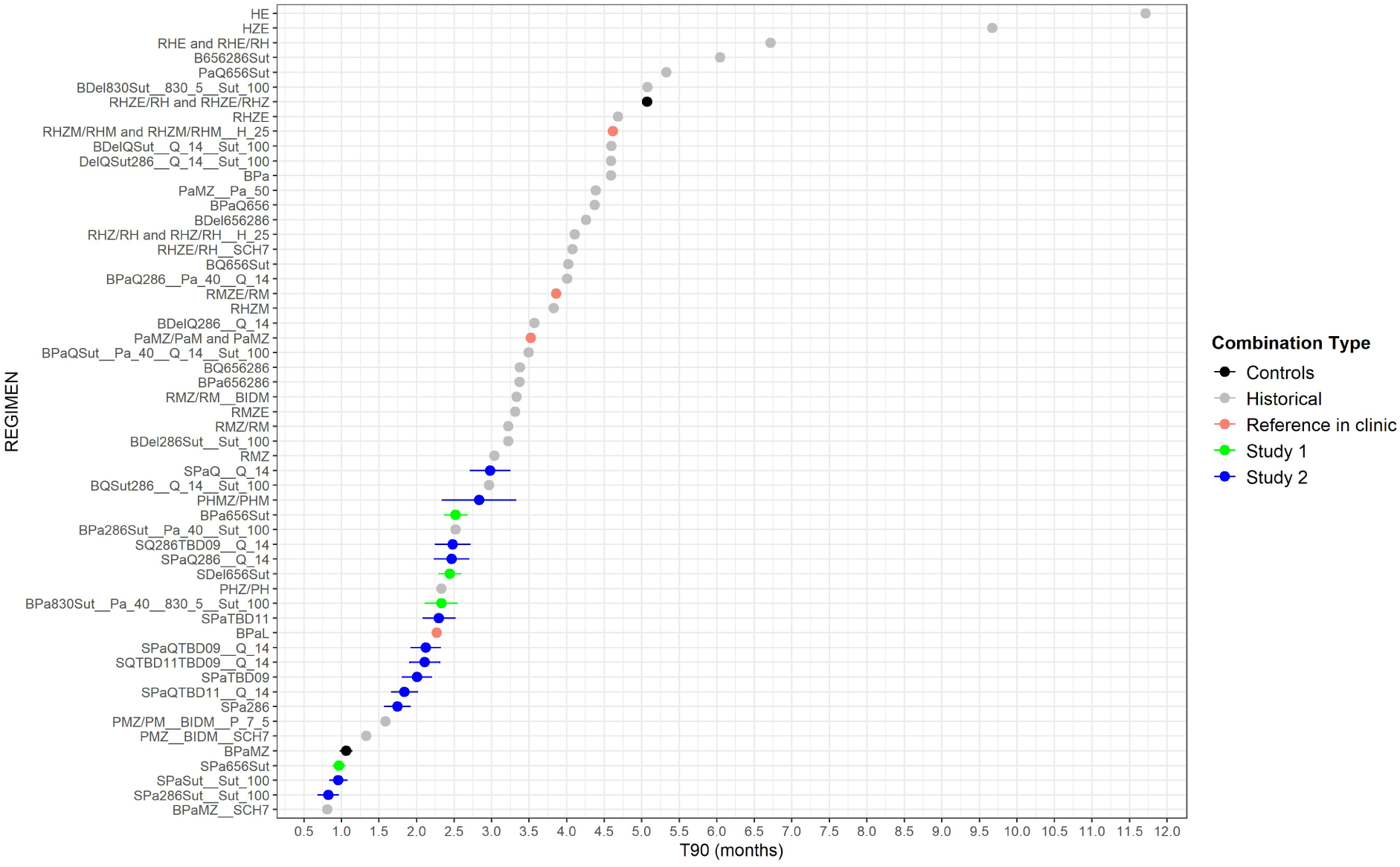
Ranking of population T90 values with 95% confidence intervals for all TB drug combinations from test and historical datasets (historical data + Study 1 and Study 2). BIDM indicates that Moxifloxacin was dosed twice in a day. SCH7 indicates that the regimen was dosed 7/7 days a week.

All combinations with T90s equal to or less than 2.1 months included S rather than B. When T90s are directly compared between BPa656Sut (2.5 months) and SPa656Sut (1 month) a reduction in T90 of around 1.5 months is observed when S is substituted for B (**Table 5** and **Figure 2**). Comparison of SPa656Sut and SDel656Sut indicated a greater efficacy contribution of Pa than Del in this combination, with a T90 difference of about 1.5 months between the two combinations (**Table 5** and **Figure 2**). Finally, direct comparison of equivalent combinations including each of the two leucyl-tRNA synthetase (LeuRS) inhibitors, 656 and 830, indicated that they contributed efficacy similarly to each other when added to BPaSut (2.5 vs 2.3 months) (**Table 5 and Figure 2, 6 and S3**).

### A treatment-shortening contribution of Pa, Sut, TBD09, Q and TBD11 or 286 in S containing combinations

Comparison of the T90s for 3-drug vs 4-drug combinations (i.e., where a 4^th^ drug was added) demonstrated that addition of Pa or TBD11 to SQTBD09, resulted in a decrease of time to cure 90% mice of at least 1 month, i.e. 100% relapse was observed after 3 months’ treatment for SQTBD09 whereas the T90 for both SQPaTBD09 and SQTBD09TBD11 was 2.1 months. Similarly, the presence of Sut in SPa286Sut (0.8 vs 1.7 months) or TBD09 in SPaQTBD09 (2.1 vs 3 months), Q or TBD11 in SPaQTBD11(1.8 vs 2.3 or 3 months), or 286 in SPaQ286 (2.5 vs 3 months) significantly decreased the time needed to cure 90% mice **(Table 5**, **Figures 4 and 5 and Table S3**). Conversely, the removal of Q from SPaQ286 (2.5 vs 1.7 months) resulted in a significant decrease in T90 (**Table 5 and Figure 5 Table S3)**, corresponding to an improvement in time to cure of 0.72 months. The reasons for this result are unknown.

**Figure 4.**
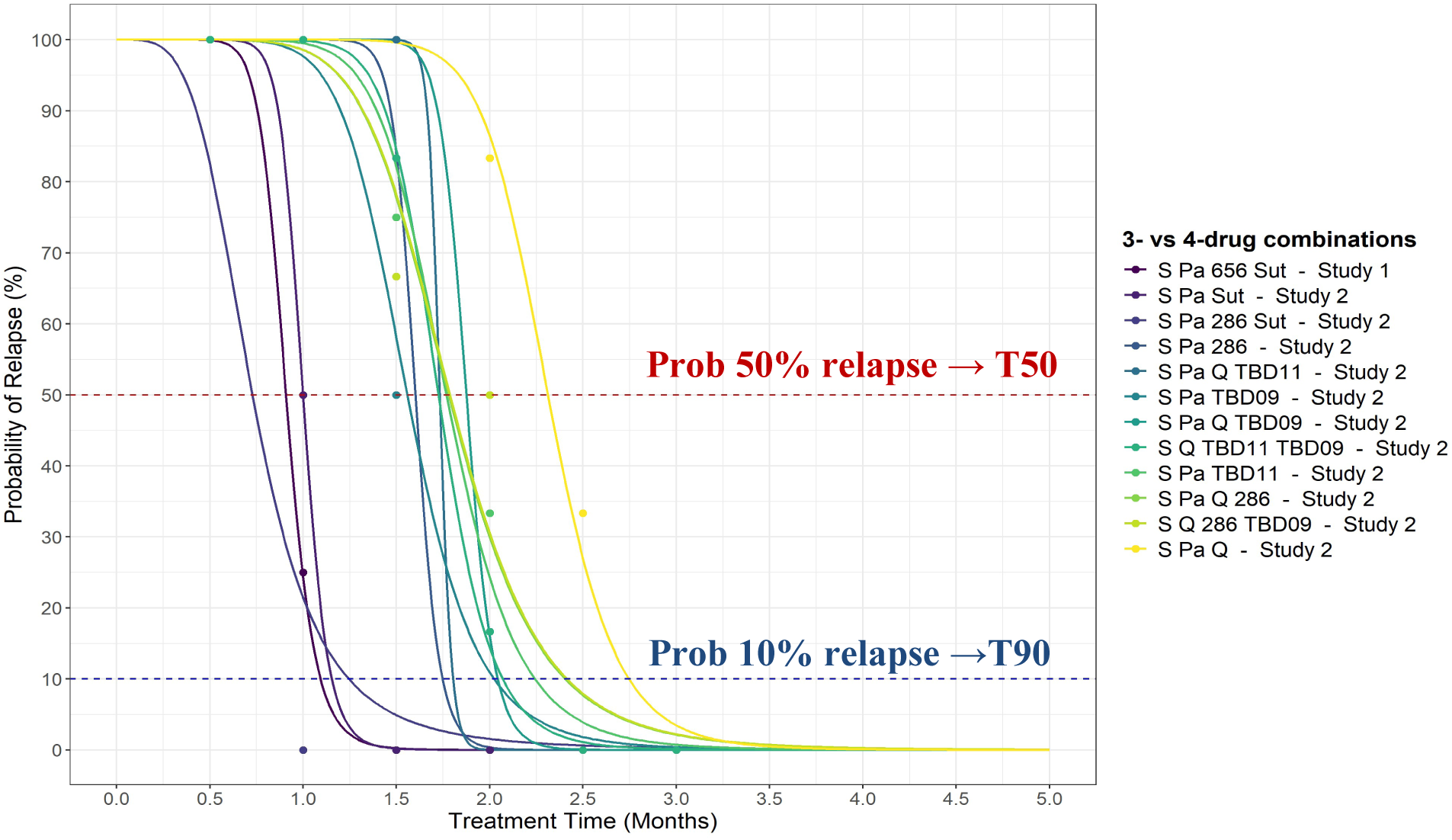
Probability of Relapse with Treatment Duration for BALB/c Mice treated with combinations tested in Study 1 and Study 2 studies. Comparison of 3- vs 4- drug combinations.

**Figure 5.**
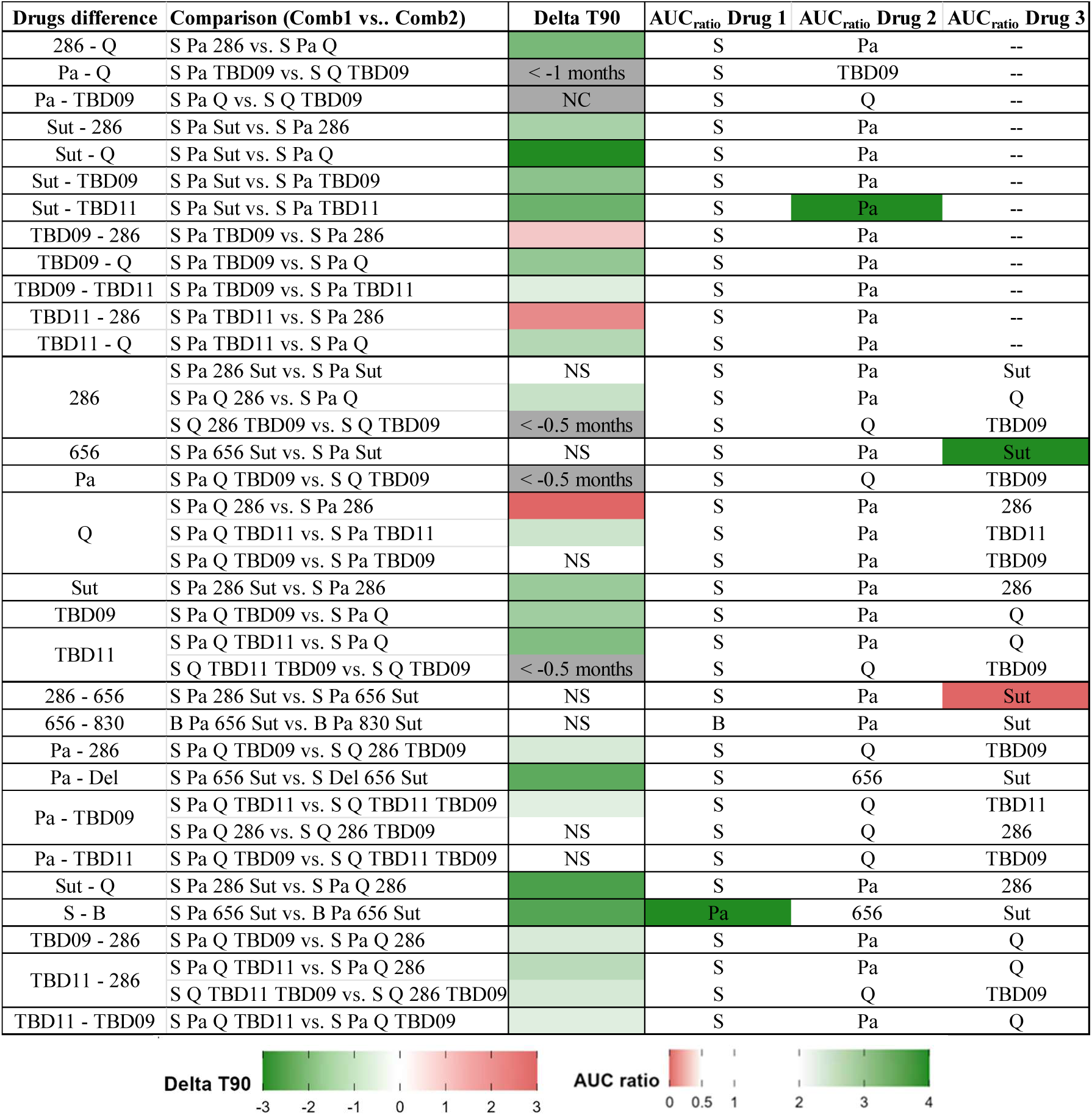
Statistical comparison of T90 and AUCratio between combinations differing by one drug. Green (Delta T90 < 0) and red (Delta T90 > 0) indicate statistically significant differences (p < 0.05) in T90 between Combination 1 and Combination 2. Green (AUCratio >2; AUC_0-24_ DrugCombination1 > 2×AUC_0-24_ DrugCombination2) and red (AUCratio <0.5; AUC_0-24_ DrugCombination1 < 0.5×AUC_0-24_ DrugCombination2) indicate statistically significant differences in AUCratio. Grey cells indicate not available delta T90 due to not computable T90 values (not computable T90 values are assumed to be > 3 months). NS means “Not significant” (p > 0.05). NC means “Not Computable” T90 value. 830 and 656 are considered as equivalent compounds.

The comparative efficacy contributions of TBD11 versus 286 and TBD09 versus Sut, to both 4- and 3-drug combinations, were also assessed. In particular, the replacement of TBD09 by Sut in the 3-drug combination SPaX resulted in a significant reduction in T90 of about 1 month. The observed effect on T90 of replacement of 286 by TBD11 was combination-specific with an increased T90 in the context of SPaX (1.7 vs 2.3 months) and a decreased T90 in the context of SPaQX (2.5 vs 1.8 months) and SQXTBD09 (2.5 vs 2.1 months) (**Table 5 and figure 5 Table S3**).

### RMM plasma exposures were within 2-fold of their clinical targets for most combination components

The doses of each drug used in the RMM studies had been selected based on previous data (Sordello et al, 2026) or existing nonclinical PK data available to PAN-TB, with the aim of achieving steady-state area under the concentration time curve from 0-24 hours (AUC_0-24_) for each drug, across all test combinations, within 2-fold of observed or targeted clinical plasma concentrations (**Table 6**).

**Table 6.** Doses utilized in RMM studies to reach clinical target plasma exposures : Numerical distribution of plasma AUC_0-24_ (*AUC_0-24MIC_ (h) for B, S and Sut) from popPK analysis and corresponding target value and 2-fold range.

| Compounds<br>(Blood to plasma ratio) | Dose selected<br>for RMM<br>studies (mg/kg) | Mouse<br>AUC<sub>0-24</sub><br>(µg*<sup>h</sup>/mL) | Target [2-fold<br>range] human<br>plasma AUC<sub>0-24</sub><br>(µg*<sup>h</sup>/mL) | Reference for<br>human target<br>plasma AUC<sub>0-24</sub> |
| --- | --- | --- | --- | --- |
| Bedaquiline [B]* (0.60) | 25 | 412*<br>[288-434] | 403*<br>[202-807] | NDA 204384 |
| Pretomanid [Pa] (1.71) | 40 | 59<br>[45-77] | 52<br>[26-104] | NDA 212862 |
| Delamanid [Del] (0.48) | 2 | 13.29<br>[12.76-14.63] | 5.90<br>[2.95-11.80] | Provided by Otsuka |
| Sutezolid [Sut]* (1.57) | 100 | 131*<br>[58-145] | 188*<br>[94-376] | (Wallis et al., 2014) |
| Quabodepistat [Q] (0.64) | 14 | 12.18<br>[9.69-13.08] | 2.50<br>[1.25-5.00] | Dawson R et al<br>2025 |
| GSK2556286 [286] (1.04) | 35 | 23.03<br>[19.65-26.51] | 22.50 (predicted)<br>[11.25-45.00] | provided by GSK |
| GSK3211830 [830] (2.09) | 5 | 1.34<br>[1.30-1.39] | 6.50 (predicted)<br>[3.25-13.00] | provided by GSK |
| GSK3036656 [656] (2.04) | 8 | 5.20<br>[4.71-6.02] | 6.50<br>[3.25-13.00] | (Tenero et al 2019) |
| TBD09 (0.66) | 200 | 578<br>[526-640] | 224 (predicted)<br>[112-448] | provided by GMRI |
| TBD11 (1.07) | 5 | 10.0<br>[5.9-13.0] | 5.6 (predicted)<br>[2.8-11.2] | provided by GMRI |
| Sorfequiline (TBAJ-876) [S]*<br>(0.80) | 6.25 | 773*<br>[696-884] | 1307*<br>[653-2613] | Provided by<br>TBalliance |

To evaluate the PK exposures achieved during the RMM efficacy studies for each drug across the test combinations, a population PK (popPK) analysis was performed using sparse-sampled drug concentration data obtained after 8 weeks’ dosing during the RMM studies, together with rich PK data generated through combination PK studies. The combination PK studies were performed in infected BALB/c mice in parallel with the RMM studies, for the corresponding 4-drug combinations with compounds administered at the selected doses.

Individual RMM drug exposure distributions across all tested combinations are shown in **Figures S2-S10**. For most of the tested compounds, i.e., 286, 656, B, Pa, S, Sut and TBD11 the median observed plasma AUC_0-24_ values achieved after 8 weeks’ dosing during the RMM studies were within 2-fold of their clinical targets (**Figure 6**, **Table 6**). For Del (only one combination) and TBD09 the median observed plasma AUC_0-24_ values were slightly more than 2-fold higher than the target (**Figure 6 and Figure S9**). The 830 RMM exposure (only one combination) was more than 2-fold lower than the clinical target. Finally, the median AUC_0-24_ observed for Q in the combinations tested here was 4-fold above the clinical target (**Figure 6 and Figure S6).** Because Q had previously demonstrated substantial regimen-dependent exposure variability in mice, the dose of Q for use in these studies was selected with the aim of maintaining the clinical target across regimens. The consistent (i.e less regimen-dependent) exposures observed in these studies therefore led to high median exposure. For all drugs studied herein, formal PK/PD evaluations performed in the context of combinations of interest will be needed to support a full understanding of exposure-response relationships for individual drugs within specific combinations.

**Figure 6.**
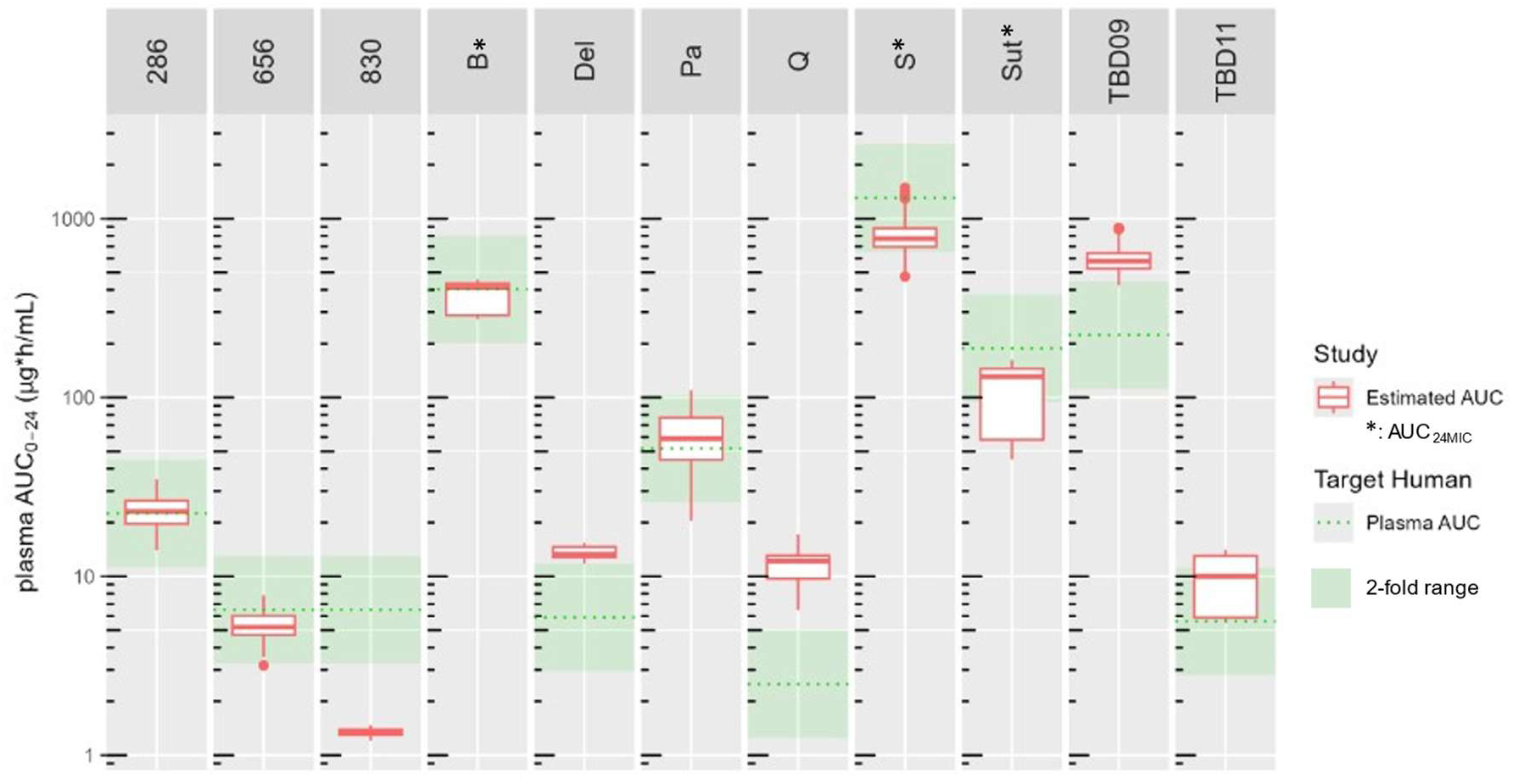
Distribution of plasma AUC_0-24_ (AUC_24MIC_ for B, Sut and S) from popPK analysis versus clinical target. AUC_0-24_, are under the concentration time curve from 0 to 24 hours; AUC_0-24MIC_, AUC_0-24_-to-MIC_90_ ratio;

## DISCUSSION

Our previous work utilizing this BALB/c RMM workflow, within a modeling-based framework, demonstrated the treatment-shortening potential of 15 first-generation PAN-TB 4-drug combinations, including 3 (BPa830Sut, BPa286Sut and BQSut286) that cured 90% of mice in less than 3 months. Here, using combinations that included next-generation PAN-TB candidates with the potential to demonstrate superior potency, safety and PK properties compared to first-generation drugs, we identified 14 novel 3- or 4-drug combinations with pan-TB potential and shorter time to cure than PHMZ. PHMZ, evaluated here in a BALB/c RMM for the first time, has demonstrated clinical treatment-shortening compared to RHZE/RH and also cured 90% of mice faster than RHZE/RH in this animal model.

Among the next-generation combinations, SPa286Sut, SPaSut and SPa656Sut cured 90% of mice with 1 month of less treatment - at least as fast as BPaMZ. Three additional combinations (SPa286, SPaQTBD11 and SPaTBD09) sterilized mice with less than 2 months’ treatment which is faster than BPaL in this model (T90 of 2.3 months). All of these novel combinations sterilized mice more rapidly than BPaQSut, which was recently evaluated with BDelQSut in Gates MRI-TBD06-201 PAN-TB trial (https://ClinicalTrials.gov ID NCT05971602). This trial concluded that although both regimens demonstrated strong bactericidal activity by the end of treatment when administered for 4 months, neither showed sufficient evidence for being able to treat TB in 3 months or less based on 2- and 3-month sputum culture conversion rates and TB recurrences after treatment completion. When assessed by 12-month post-randomization unfavorable outcome rates, the 4-month PBQS regimen performed similarly to 6-month HRZE (Int J Tuberc Lung Dis 2025: 29 (11 Suppl 1): S1 – S963). Our previous work [12] indicated that results from this mouse model were consistent with these clinical results: The population derived T90 in mice for BPaQSut exceeded 3 months (having a value of 3.5 months) but was significantly shorter than that of RHZE/RH (5.1 months) (**Figure 7).**

**Figure 7.**
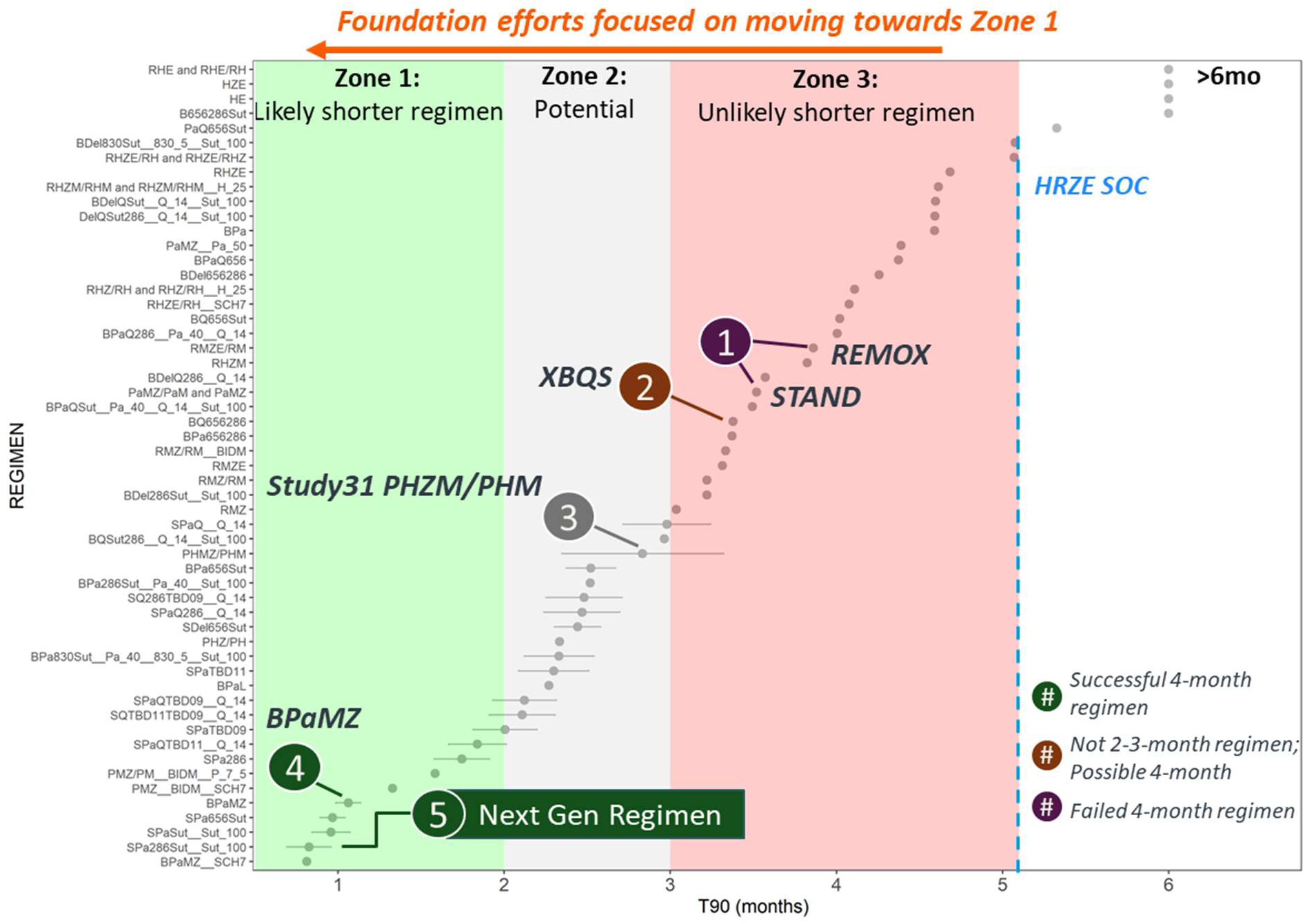
Ranking of population T90 values with 95% confidence intervals for historical and test regimens, indicating clinical benchmarks.

All of the novel combinations that cured 90% of mice in less than 2 months include S as the diarylquinolone, rather than B. In S/BPa656Sut and in S/BPa830/656Sut regimen pairs, we were able to compare the contributions of B versus S (assuming that 830 and 656 had equivalent contributions within S/BPa830/656Sut). Replacing B with S at this dose in these regimens resulted in an average reduction in time to cure of 1.5 months. These results are in line with previous findings from BALB/c RMM studies, demonstrating the superior treatment-shortening contribution of S versus B; for example, SPaOxa (TBI-223) cured BALB/c mice faster than BPaOxa [17]. The dose of S utilized here resulted in an average exposure in mice similar to that achieved following administration of 50 mg S to patients in a Phase II trial conducted by TB Alliance NC-009 (https://ClinicalTrials.gov ID NCT06058299). Clinical data have indicated the potential to safely administer higher doses of S to patients, to enable evaluation of efficacy at higher exposures - for example, 100 mg S was administered in the NC-009 Phase II trial and was shown to be well-tolerated and provided faster time to sputum culture conversion than B when combined with PaL. Future studies in BALB/c RMM and other nonclinical models will enable probing the exposure-response relationship of S within novel drug combinations of most interest.

In addition to S, Pa was present in all next generation combinations with T90s less than 2 months, while Del was included in only one. When comparing the sterilizing contribution of Del versus Pa in the two regimens in which the other 3 drugs were the same (SPa656Sut versus SDel656Sut), the contribution of Pa was greater than that of Del in this model. There is evidence for inhibition of synthesis of cell wall components as part of the complex mode of action of the nitroimidazole class. The novel drug candidate Q inhibits synthesis of arabinogalactan and is also classified as a cell wall inhibitor. In our previous work, by comparing 4-drug combinations where one drug differed, we identified an apparent treatment-shortening contribution of Q that was similar to that of the nitroimidazoles Pa or Del, within specific B-containing combinations (BQSut286 vs BDel286Sut (2.9 vs 3.2 months) or BQ656286 vs BPa656286 (3.4 vs 3.4 months) or BQSut286 vs BPa286Sut (2.9 vs 2.5 months). In that context, it is of interest that the present study demonstrates similar contributions for Q and Pa in SXTBD09 and indicates treatment-shortening by Q when added to SPaTBD11 (T90s of 2.3 months versus 1.84 months). To our knowledge, this is the first formal demonstration of a sterilizing contribution by a DprE1 inhibitor to an S-containing regimen in this mouse model. Moreover, based on this data, SPaQTB11 represents a powerfully sterilizing drug combination that is oxazolidinone-free – providing an additional 4-drug combination option with pan-TB potential, beyond those based on a diarylquinoline-oxazolidinone core.

With the exception of the above-described SPaQTBD11 and 3-drug combination SPa286, an oxazolidinone is present in all evaluated combinations with T90s of less than 2 months. The combinations tested here included two oxazolidinones: Sut, which has demonstrated efficacy in clinical trials (https://ClinicalTrials.gov ID NCT07094932; [20]) and the newer drug candidate TBD09, which is now entering clinical Phase II and has potential for improved PK and safety compared to Sut (https://ClinicalTrials.gov ID NCT05971602). Although the treatment-shortening contribution of TBD09 was inferior to that of Sut within the context of SPaX (SPaSut T90 0.96 months vs SPaTBD09 2 months), TBD09 demonstrated a treatment-shortening contribution within SPaQTBD09 – a promising result for this next generation oxazolidinone.

Beyond S, nitroimidazoles and Q, the oxaborole leucyl tRNA synthetase inhibitor, 656 (ganfeborole), features within one of the combinations (SPa656Sut) that cures 90% mice in less than one month. This advanced clinical development candidate is of the same chemical class and shares a mechanism of action with 830, which was evaluated within other combinations in our previous work. Herein we made a direct comparison of regimens that include either 656 (ganfeborole) or 830, and see that they contribute similarly to each other, in the context of B. The clinical EBA trial data for ganfeborole [21, 22] illustrates the clinical efficacy potential of this oxaborole candidate drug, which is under evaluation for efficacy and safety within Phase II combination trials, including those conducted by UNITE-4TB. The preclinical efficacy data presented here further highlights leucyl tRNA synthetase as an exciting novel drug target for both DS- and DR-TB.

Finally, both evaluated cholesterol metabolism inhibitors, 286 and TBD11, are present in the drug combinations with T90s of less than 2 months. Several nonclinical studies have illustrated the potential of compounds with this new mechanism, utilizing 286 [23, 24, 12]. It is promising that TBD11, a newer molecule available to replace the now-discontinued 286, was able to contribute sterilizing activity similarly to 286 within combinations studied (SPaQ, SQTBD09). Further evaluation of TBD11 within the context of additional drug combinations is warranted as this molecule advances in clinical trials (https://ClinicalTrials.gov ID NCT06707142).

Overall, the fastest-to-cure 4-drug combinations identified in this study were SPa286Sut and SPa656Sut. One of these combinations includes the discontinued compound 286. However, we demonstrated that TBD11 can replace 286, offering a similar sterilizing contribution, thus enabling development of a regimen including the novel cholesterol metabolism inhibitor. The replacement of Sut with the potentially safer and lower clearance TBD09 may also offer benefits for a broad pan-TB patient population. The identification of these rapidly curative 4-drug combinations is exciting for the TB field. While ganfeborole-containing combinations are under study by the UNITE-4TB consortium, the PAN-TB consortium recently endorsed plans for further clinical and non-clinical evaluation of the SPaTBD11TBD09 combination, based in part on these promising BALB/c RMM data.

The BALB/c RMM model and methodology used here has limitations. These include the lack of advanced disease pathology in the BALB/c mouse, a hallmark of hard-to-treat disease in humans. For this reason, the PAN-TB consortium considers the presented evaluations as a first step in nonclinical combination evaluation and prioritization. However, it is notable that, despite its shortcomings, this mouse model is able to correctly rank-order T90s for several drug combinations, where clinical data is available for comparison. For example, BPaMZ and PHMZ demonstrated shorter T90s than RHZE/RH in this model, as demonstrated in a clinical trial for PHMZ [18] and indicated in a clinical trial for BPaMZ [19]. Additional nonclinical and clinical data – including evaluations in mouse TB models with more human-like pathology, evaluations of lesion penetration and activity of drugs and combinations against *Mtb* in caseum – need to be performed to better assess potential efficacy in the clinic for the promising next generation combinations identified here. However, demonstrated T90s for novel pan-TB combinations that are shorter than those for BPaQS, BDelQS, PHMZ, BPaL and BPaMZ are a promising first step.

A further limitation of the work is the lack of a complete understanding of exposure-response (PK/PD) for the drugs tested here within the context of the evaluated combinations. Although the exposures for most compounds were within 2-fold of the (clinical) AUC_0-24_ targets across most combinations, this was not always the case, challenging our interpretation of the efficacy data. For example, although a sterilizing contribution of Q was evident within the context of SPaTBD11, the addition of Q to SPa286 resulted in an increased T90, i.e. apparently reducing the sterilizing activity of SPa286 while increasing the sterilizing efficacy of SPaTBD11 despite 286 and TBD11 sharing the same mechanism of action and without any significant changes in AUC_0-24_ of the partner drugs. Further work is required to understand this apparently paradoxical result and these results highlight the challenges inherent in interpretating combination efficacy data in the absence of extensive information on pharmacological interactions or detailed PK/PD.A further example of the limitation of this dataset to precisely interpretation efficacy contributions across multiple 4-drug regimens is presented by Q: Because Q had demonstrated regimen-dependent exposure variability in mice, when evaluated in the context of combinations tested in our previous work [12], a higher Q dose was selected for use here, to ensure maintenance of the clinical target across the present test regimens. In fact, we observed consistent exposures and therefore a median AUC_0-24_ 4-fold above the target. In the absence of detailed PK/PD in mice we cannot determine whether the contribution to combination efficacy was greater for Q at higher than lower exposures. However, in clinical studies, similar efficacy was observed after dosing either as monotherapy (in a bactericidal activity study) or as part of the 3-drug combination Q+B+Del, with the 30 mg daily dose that is equivalent to the dose used in this study and the 90 mg daily dose (which achieved exposures higher than observed in this study).This suggests a potential efficacy plateau at the 30 mg dose and relevant exposure achieved in humans. Therefore, the higher-than clinical target Q exposure observed here in mice may not have resulted in an increased efficacy contribution for Q in the tested combinations.

Further to these limitations, translation between mouse and clinical settings relied on matching plasma AUC_0-24_, while other potentially relevant pharmacokinetic drivers were not considered in these investigations. It will be of great interest to further probe the efficacy of specific combinations of interest and the contributions of specific agents across exposures, to inform predictive modeling approaches towards clinical dose selection.

Successful development of an all-oral 4-drug combination with pan-TB potential, for example SPaTBD11TBD09, could bring positive impact to both patients and healthcare systems. Further benefits are likely to come from additional innovations in TB treatment such as administration of drugs as Long Acting Injectables (LAIs). Administration of a 4-drug combination for a short period of time followed by some or all of those drugs as a single encounter LAIs could reduce the burden on patients and caregivers and decrease potential for missed doses. LAI formulation development is underway for several PAN-TB drugs and drug candidates. Further work is needed to better assess the impact of such administration versus an all-oral combination for the SPaTBD11TBD09 and other promising regimens, considering anticipated differences in PK profiles between oral and LAI administrations. However, the potential to further shorten and simplify pan-TB treatments may offer a transformative step-change in TB treatment, worthy of exploration. In summary, this work demonstrates that the next-generation TB drug combinations evaluated here can achieve more rapid sterilization than clinically available regimens in this preclinical model, at clinically-relevant exposures. These results highlight their promise as shorter, safer pan-TB regimens with potential to improve outcomes for patients and healthcare systems.

## MATERIAL AND METHODS

### Animals and ethics

All mouse experiments were carried out at the Evotec France SAS animal facility. This facility is accredited by the French Ministry of Agriculture and by the Association for Assessment and Accreditation of Laboratory Animal Care International (AAALAC). All studies were performed under the European Communities Council Directive (2010/063/EU) for the care and use of laboratory animals and approved by local Ethical Committee CEPAL: CE 029 and authorized by the French Ministry of Education, Advanced Studies, and Research. Six-week-old female BALB/cJRj mice from Janvier Laboratories were group housed in bioconfined cages (Isocage, Tecniplast®) under a 12h light:12h dark with free access to filtered water and a standard rodent diet (AO4C, Safe, France). An ambient temperature of 22 ± 2°C, a relative humidity of 55 ± 10 %, and a negative pressure of -20Pa were maintained throughout the study. All mice were allowed to acclimatize to their new environment for at least 5 days prior to the start of the study.

### Drug formulations and dosing strategies

Drugs were acquired or provided by PAN-TB consortium members, and formulations prepared for dosing as follows at the selected doses. B (J&J IM) was formulated in 20% 2-hydroxypropy-β-cyclodextrin pH 3 for dosing at 25 mg/kg; Pa (TB Alliance) was formulated in 10% Hydroxy-propyl-beta-cyclodextrin and 2% soy lecithin for dosing at 100 mg/kg for the BPaMZ control group and at 40 mg/kg for dosing in the PAN-TB combinations. M (LTK Laboratories) and Z (Sigma) were co-formulated in water for dosing at 100 mg/kg and 150 mg/kg, respectively. Del (Otsuka) and Q (Otsuka) were formulated in 5% Arabic gum for dosing at 2 and 14 mg/kg, respectively. 286, 656, and 830 (GSK) were formulated in 1% methylcellulose solution for dosing at 35, 8, and 5 mg/kg, respectively. Sut (TB Alliance) was formulated in 0.5% methylcellulose and 5% PEG200 for dosing at 100 mg/kg. P (Sigma) was formulated in water for dosing at 10 mg/kg. H (Sigma) was formulated in water for a dosing at 10 mg/kg. TBAJ-876 (TB Alliance) was formulated in 20% (w/v) hydroxypropyl-beta-cyclodextrin (HP-β-CD) and 0.1% (w/v) Tween 20 in citrate buffer (50 mM, pH=3.3 [±0.3]) for dosing at 6.25 mg/kg. TBD09 (formerly MK-7762) (GMRI) was formulated in 10% Tween 80/40% PEG400:50% water vehicle for dosing at 200 mg/kg. TBD11 (GMRI) was formulated as a suspension in 0.5% methylcellulose/0.5% Tween 80 (MC/T) solution for dosing at 5 mg/kg. For each PAN-TB combination, each drug was dosed individually with an interval of 2 hours between each drug. The order of dosing for each drug in each combination was based on the half-life of each drug where drugs with longest half-lives were administered first and drugs with shorter half-lives were given later in the day. The drug dosing order for each combination is indicated in the name of the combination i.e. the drug listed first was dosed first, and so on. For the BPaMZ comparator, B was administered first, followed by Pa and finally MZ, dosed as a co-formulation. For the PHMZ comparator, P and H were dosed individually, followed by co-formulated PHMZ during the first 2 months and then followed by 1 month of PHM dosing MZ where P, H and M were individually administered. The same 2h interval was applied between 2 administrations for the comparators as for the test combinations.

### Relapsing mouse model

The test combinations were evaluated in two studies. The first (study 1) is named RMM2023_2 in the C-PATH APEX database: https://c-path.org/tools-platforms/tb-apex, and the second (study 2) is named RMM2024_1 in the C-PATH APEX database: https://c-path.org/tools-platforms/tb-apex). BPaMZ and PHMZ comparators were included in each Study. Both studies included 4 to 6 mice per group, per time point, allocated following a similar approach to that reported previously [25] and described in **Table S1**. *M.tb* H37Rv stock solution was prepared at exponential growth phase in 7H9 medium / 10% OADC (oleic acid-albumin dextrose-catalase) / 15% glycerol. At Day -14, female BALB/c mice were anaesthetized with 2.5% isoflurane in 97.5% oxygen and were intranasally infected with 50µL *M.tb* H37Rv at an inoculum level of 4.5 Log_10_ CFU/mouse. The bacterial burdens in the lung 1 day and 14 days post infection were 5.3 ± 0.13 and 7.5 ± 0.2 Log_10_ CFU/g for Study 1 and 5.5 ± 3 and 7.3 ± 0.26 Log_10_ CFU/g for Study 2. Treatment started 14 days post infection, designated Day 0. All drug combinations were dosed once daily by oral gavage for 5 days a week (omitting the weekend) for 2, 4, 6, 8, 10, or 12 weeks. For analysis of mouse lung bacterial burden at the end of each designated treatment period, mice were sacrificed 24h post last dosing. For relapse assessment, mice were sacrificed 12 weeks after the end of each designated treatment period. Following sacrifice, lungs were collected, weighed and lung samples homogenized and plated undiluted, or serially diluted on 7H11-OADC + 0.4% activated charcoal. Plates were incubated at 37°C for 6 weeks for CFU quantification.

### Pharmacokinetics

Combination PK studies were conducted, before the RMM studies, in female BALB/c mice, intranasally infected with *M.tb* H37Rv in the same manner as for the RMM studies (see RMM section of Materials and Methods). Beginning 14 days post-infection (Day 0) 24 mice per group were treated with each test combination via oral gavage either once or once daily for 5 days. The 5-day dosing period was selected based on the short terminal half-lives of the drugs (**Table S3**), suggesting at least 80% of the steady-state is reached after 5 days treatment duration (5 days > 5 fold T_1/2_). The only exceptions are the M3 Soferquiline and the M2 Bedaquiline metabolites, exhibiting a longer half-life. For both, independent PK simulations show that after 5 days repeated administration, > 80% of the steady-state concentration obtained in the RMM efficacy study, is reached. In addition, it is assumed that the late sparse PK samples in the RMM study (Week 8) provide enough information to correctly estimate the PK parameters of these entities. Doses in ancillary PK study were selected based on historical data from the partners. Exposures observed in the ancillary PK studies were compared with the clinical plasma AUC and used to tailor the RMM studies doses, as shown in **Table 6**. The order and spacing of administrations were performed as for the RMM studies (see Dose Formulations and Dosing Strategies). Blood samples were collected from the tail vein on days 1 (after single dosing) and 5 (after once daily repeat administration) at 0.5, 2.5, 5.5, 7.5, 10, 12, 14, 24, 36, 29, and 31 hours post administration of the first drug. Additionally, for each combination, plasma and blood intra-cardiac concentrations were assessed on Day 1 and Day 5 at 2.5, 7.5, 14, and 31 hours post-first compound dosing. During RMM studies, on the 5^th^ day of the 8^th^ week of treatment, blood was collected from the tail vein at 0.5, 2.5, 5.5, 7.5, 9 and 24 hours post-first compound dosing. In all cases, after blood processing, each compound, including the bedaquiline M2 (desmethyl) metabolite (B-M2), Sorfequiline and its metabolite M3 (S-M3) and the active (sulfoxide) metabolite of sutezolid (PNU-101603) were quantified by liquid chromatography tandem mass spectrometry (LC-MS/MS). Bioanalytical methods, samples handling, as well as MS conditions are described in supplemental data. All PK assessments were conducted in blood. Blood and plasma concentrations determined through the ancillary PK studies were used to calculate mouse geometric mean Blood-to-Plasma (BP) ratios (**Table 6**) for each drug. These values were used to facilitate translation of blood to plasma exposure values for head-to-head comparisons between observed mouse and clinical target exposures (see Population PK Modelling).

### Population PK Modelling

For each compound tested across the PAN-TB combinations, a population pharmacokinetic (popPK) model was developed to describe the mouse PK profile. After identifying each specific model, post-hoc estimates were used to calculate total exposures, i.e. AUC_0-24_, accounting for the variability observed across animals in terms of summary statistics (median, 5^th^, 25^th^, 75^th^ and 95^th^ percentiles). Blood concentration data collected during the RMMs were pooled with those collected in the ancillary PK studies to ensure a sufficient number of observations required for a robust model development and parameters estimation (more details about model selection criteria are provided in the supplemental material). For S, B and Sut their main active metabolites, S-M3, B-M2 and PNU-101603 respectively, were included in the model description to derive the overall exposure. As the parent drug and corresponding metabolite contribute to efficacy in mice and human at different relative levels [26, 27, 28, 29], the active moiety approach was adopted to compare the relevance of preclinical exposure values to clinical exposures, considering the plasma AUC_0-24_/MIC_90_ ratios (AUC_24MIC_) in both human and mouse:

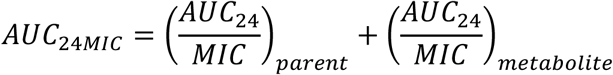

To enable comparisons of mouse exposures against clinical plasma targets, simulated blood concentrations were converted to plasma values using the compound-specific geometric mean Blood-to-Plasma (BP) ratio (**Table 6**). The area under the curve, during the 24 hours post drug administration (AUC_0-24_, ng·h/mL) was determined, assuming a dense simulation time grid (i.e., 0.1 h sampling frequency) which enabled robust determination of the total exposure in plasma. PopPK modelling was performed with Monolix Suite^®^ (Build 2024R1, Lixoft) running on a Windows 11, 64bit computer. All models were identified using an SAEM estimation algorithm. All analyses for the extrapolation of exposure metrics were performed using R Statistical Software (v4.4.2; R Core Team 2024).

### Logistic Emax Model

A logistic Emax model was applied, as previously described [25] using data obtained from Study 1 and Study 2 plus a historical dataset. The historical dataset consisted of data utilized in previous modeling efforts [30] as well as other literature data [31, 32, 33] and historical data generated at Evotec. The Evotec internal studies assessed efficacy of RHZ/RH, BPaL and PaMZ (where L stands for Linezolid drug), dosed 5/7 or 7/7 days per week in a BALB/c RMM and data from these studies is available via the C-PATH APEX platform (https://c-path.org/tools-platforms/tb-apex).

Data from the same treatment collected from different studies (i.e. within the historical and present datasets) were pooled together. For each combination i and study k, γ and T50 parameters were estimated and subsequently used to calculate the related T90, according to the following formula:

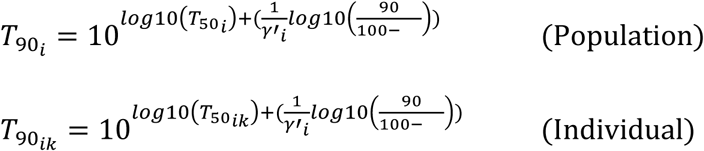

Population parameters (e.g. T50_i_ and T90_i_) enable characterization of the average time to 50% or 90% cure for a particular regimen shared across studies independently from study-related factors (as for example the starting inoculum), while the individual parameters (e.g. T50_ik_ and T90_ik_) provide the average time to 50% or 90% cure for each specific study. Notably, modeling could not be performed for drug combinations where relapse was observed in 100% of mice at the last timepoint tested.

Since the model estimation algorithm was not able to converge when estimating T50 and γ parameters for historical, Study 1 and Study 2 combinations together due to the high number of parameters to be estimated (more than 100), a two-step identifiability process was considered: T50 and γ were firstly estimated for historical combinations only, thus excluding Study 1 and Study 2; then, the model was identified by estimating T50 and γ for Study 1 and Study 2 combinations, while fixing parameters related to the other combinations to the corresponding values obtained in the previous model iteration.

Once all these values were computed for each combination, a specific ranking was drawn up in order to compare the performance of the novel combinations with respect to historical ones and controls (i.e. BPaMZ and PHMZ/PHM).

To obtain estimates of model precision and support comparison of the various regimens, confidence intervals (CI) for T90 parameters were computed by deriving an approximation of T90 standard error from T50 and γ estimates and then computing 2.5^th^ and 97.5^th^ percentiles by assuming normally-distributed data, as previously described [25]

The logistic Emax model was developed using NONMEM® 7.5.1 and data handling was performed using SAS® version 9.4 and R Statistical Software (v4.4.2; R Core Team 2024).

### Statistical analysis

Statistical analysis was conducted to evaluate statistically significant differences between pairwise drug combinations differing by a single component, specifically comparing four-component combinations with the related three-component combinations. Compounds “830” and “656” were treated as equivalent. The analysis considered both population T90 results and exposure levels (AUC_0-24_) of the drugs common to the combinations. T90 results were compared using a Z-test, assuming normally distributed values with known standard errors. AUC comparisons were performed using a t-test with unequal variances on log-transformed data. P-values below 0.05 were considered statistically significant.

## Acknowledgements

This work was supported, by the Gates Foundation [BMGF01 INV-008993]. The conclusions and opinions expressed in this work are those of the author(s) alone and shall not be attributed to the Foundation. Under the grant conditions of the Foundation, a Creative Commons Attribution 4.0 License has already been assigned to the Author Accepted Manuscript version that might arise from this submission. Please note works submitted as a preprint have not undergone a peer review process.”

We would like to thank the Evotec BSL3 In Vivo Pharmacology, Bioanalytical and Pharmacometrics teams for their valuable support and collaboration.

We are especially grateful to Sonia Edouard, Halim Guerraoui, Adrien Dedieu, Raphaël Durant, Hafsa Aounzou, Pauline Espagnolle, Mélissa Mauzole, Constance Manso, Naomi Santin, Luane Vandenberghe, Valentine Pierre, Cindy Louvet, Mélissa Toxé, Pierre Demoux, Thomas Ladrat, Jean-Jacques Medard, Joshua Estellé, Jake Brett, Amandine Basandella, Amanda Corini, Tiphaine Mauge, Pauline Chaillé, Karim Riadhi and Alexandre Daca for their technical contribution, as well as Jeanne Jaen, Fanny Deglave and Eric Erdocain for their contributions to the bioanalytical aspects of this work.

We would like to thank the Evotec Pharmacometrics team for their valuable support and collaboration throughout this work. In particular, we are especially grateful to Giulia Calusi and Andrea Boscolo Panzin for their insightful input and contributions to the analytical aspects of this work.

## Supplemental Material

**Table S1.**
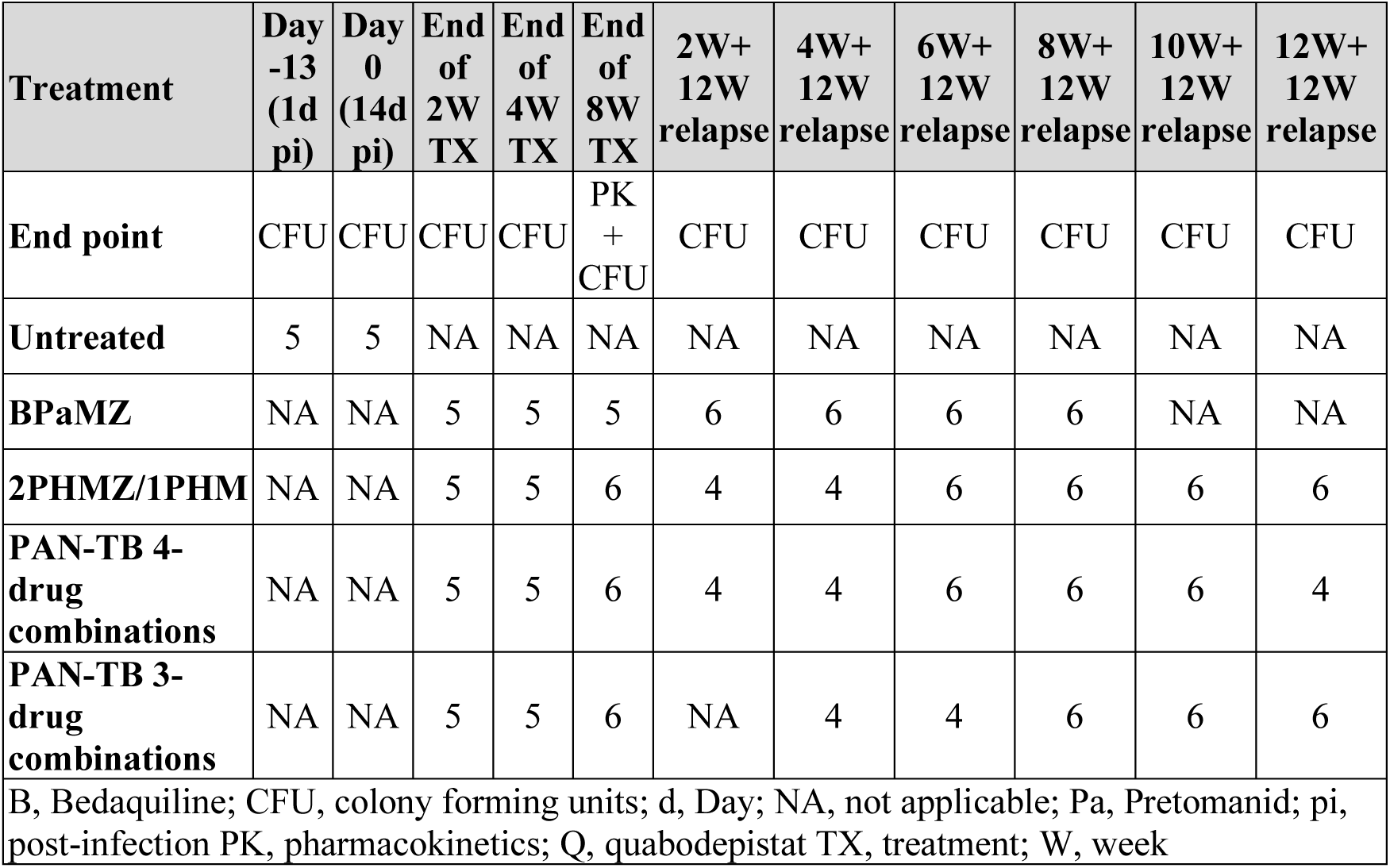
Treatment groups and number of mice per treatment group at each endpoint.

| **Treatment** | **Day**<br>**-13**<br>**(1d**<br>**pi)** | **Day**<br>**0**<br>**(14d**<br>**pi)** | **End**<br>**of**<br>**2W**<br>**TX** | **End**<br>**of**<br>**4W**<br>**TX** | **End**<br>**of**<br>**8W**<br>**TX** | **2W+**<br>**12W**<br>**relapse** | **4W+**<br>**12W**<br>**relapse** | **6W+**<br>**12W**<br>**relapse** | **8W+**<br>**12W**<br>**relapse** | **10W+**<br>**12W**<br>**relapse** | **12W+**<br>**12W**<br>**relapse** |
| --- | --- | --- | --- | --- | --- | --- | --- | --- | --- | --- | --- |
| **End point** | CFU | CFU | CFU | CFU | PK<br>+<br>CFU | CFU | CFU | CFU | CFU | CFU | CFU |
| **Untreated** | 5 | 5 | NA | NA | NA | NA | NA | NA | NA | NA | NA |
| **BPaMZ** | NA | NA | 5 | 5 | 5 | 6 | 6 | 6 | 6 | NA | NA |
| **2PHMZ/1PHM** | NA | NA | 5 | 5 | 6 | 4 | 4 | 6 | 6 | 6 | 6 |
| **PAN-TB 4-**<br>**drug**<br>**combinations** | NA | NA | 5 | 5 | 6 | 4 | 4 | 6 | 6 | 6 | 4 |
| **PAN-TB 3-**<br>**drug**<br>**combinations** | NA | NA | 5 | 5 | 6 | NA | 4 | 4 | 6 | 6 | 6 |
| B, Bedaquiline; CFU, colony forming units; d, Day; NA, not applicable; Pa, Pretomanid; pi, post-infection PK, pharmacokinetics; Q, quabodepistat TX, treatment; W, week |  |  |  |  |  |  |  |  |  |  |  |

**Table S2.** Lung bacterial burden at 2, 4 and 8 weeks of treatment in study 1 (A) and in study 2 (B): Combinations ranked based on their sterilization activity. (mean +/- sd Log_10_ CFU/ lung; (positive culture mice/total mice).

| **A_ Mean +/- SD Log<sub>10</sub> CFU/ lung (positive culture mice/total number of mice)** |  |  |  |  |  |
| --- | --- | --- | --- | --- | --- |
| **Treatment** | **Day - 13**<br>**(1d pi)** | **Day 0**<br>**(14d pi)** | **End of 2W**<br>**Treatment** | **End of 4W**<br>**Treatment** | **End of 8W**<br>**Treatment** |
| **Untreated** | 4.6 ± 0.1 (5/5) | 7.3 ± 0.2 (5/5) | - | - | - |
| **S Pa 656 Sut** | - | - | 3.5 ± 0.1 (4/4) | 0.0 ± 0.0 (0/5) | 0.0 ± 0.0 (0/6) |
| **B Pa M Z** | - | - | 3.6 ± 0.4 (5/5) | 0.4 ± 1.0 (1/5) | 0.0 ± 0.0 (0/5) |
| **B Pa 830 Sut** | - | - | 3.4 ± 0.2 (5/5) | 0.0 ± 0.0 (0/5) | 0.0 ± 0.0 (0/6) |
| **S Del 656 Sut** | - | - | 2.9 ± 0.2 (5/5) | 0.0 ± 0.0 (0/5) | 0.0 ± 0.0 (0/6) |
| **B Pa 656 Sut** | - | - | 3.8 ± 0.3 (5/5) | 0.0 ± 0.0 (0/5) | 0.0 ± 0.0 (0/6) |
| **P H M Z** | - | - | 4.7 ± 0.1 (5/5) | 2.7 ± 0.3 (5/5) | 0.0 ± 0.0 (0/6) |
| **B_ Mean +/- SD Log<sub>10</sub> CFU/ lung (positive culture mice/total number of mice)** |  |  |  |  |  |
| **Treatment** | **Day - 13**<br>**(1d pi)** | **Day 0**<br>**(14d pi)** | **End of 2W**<br>**Treatment** | **End of 4W**<br>**Treatment** | **End of 8W**<br>**Treatment** |
| **Untreated** | 5.5 ± 0.1 (5/5) | 7.3 ± 0.3 (5/5) | - | - | - |
| **S Pa 286 Sut** | - | - | 4.2 ± 0.2 (5/5) | 0.0 ± 0.0 (0/5) | 0.0 ± 0.0 (0/6) |
| **BP aMZ** | - | - | 4.5 ± 0.4 (5/5) | 0.2 ± 0.4 (1/5) | 0.0 ± 0.0 (0/5) |
| **S Pa Sut** | - | - | 4.4 ± 0.2 (5/5) | 0.4 ± 0.9 (1/5) | 0.0 ± 0.0 (0/6) |
| **S Pa 286** | - | - | 4.7 ± 0.2 (5/5) | 2.5 ± 0.2 (5/5) | 0.0 ± 0.0 (0/6) |
| **S Pa Q TBD11** | - | - | 5.6 ± 0.3 (5/5) | 2.5 ± 0.3 (5/5) | 0.0 ± 0.0 (0/4) |
| **S Pa TBD09** | - | - | 5.0 ± 0.5 (5/5) | 0.5 ± 1.1 (1/5) | 0.0 ± 0.0 (0/3) |
| **S Q TBD11 TBD09** | - | - | 4.6 ± 0.4 (4/4) | 1.3 ± 1.0 (4/5) | 0.0 ± 0.0 (0/5) |
| **S Pa Q TBD09** | - | - | 5.0 ± 0.5 (5/5) | 0.0 ± 0.0 (0/5) | 0.0 ± 0.0 (0/4) |
| **S Pa Q 286** | - | - | 5.3 ± 0.2 (5/5) | 2.1 ± 0.2 (4/4) | 0.0 ± 0.0 (0/4) |
| **S Q 286 TBD09** | - | - | 5.0 ± 0.2 (5/5) | 1.9 ± 0.1 (5/5) | 0.1 ± 0.2 (1/6) |
| **S Pa TBD11** | - | - | 4.6 ± 0.4 (5/5) | 2.4 ± 0.2 (5/5) | 0.1 ± 0.2 (1/6) |
| **PHMZ** | - | - | 5.2 ± 0.2 (5/5) | 3.3 ± 0.1 (5/5) | 0.0 ± 0.0 (0/5) |
| **S Pa Q** | - | - | 5.4 ± 0.4 (5/5) | 2.6 ± 0.2 (5/5) | 0.6 ± 0.7 (3/5) |
| **S Q TBD09** | - | - | 5.0 ± 0.3 (5/5) | 3.0 ± 0.3 (5/5) | 0.9 ± 0.6 (5/6) |

**Table S3.** Statistical comparison of T90 and AUCratio between combinations differing by one drug For each paired combination, Delta T90 = T90(Combination 1) - T90(Combination 2). For each drug, AUCratio = AUC_0-24_(Combination 1)/AUC_0-24_(Combination 2). NC means “Not Computable” T90 value. Values in brackets represent the 95% confidence interval. Asterisks mean statistical significance, specifically, * if p<0.05, ** if p<0.01 and *** if p<0.001.

| Drugs difference | Comparison (Comb 1 vs. Comb 2) | T90 Comb 1 (months) | T90 Comb 2 (months) | Delta T90 (months) | AUC<sub>ratio</sub> Drug 1 (-) | AUC<sub>ratio</sub> Drug 2 (-) | AUC<sub>ratio</sub> Drug 3 (-) |
| --- | --- | --- | --- | --- | --- | --- | --- |
| **286 vs Q** | S Pa 286 vs S Pa Q | 1.74 | 2.98 | -1.23***<br>[-1.55; -0.92] | S: 0.81*<br>[0.69; 0.94] | Pa: 1.03<br>[0.9; 1.17] | - |
| **Pa vs Q** | S Pa TBD09 vs S Q TBD09 | 2 | NC | NC | S: 0.91<br>[0.81; 1.03] | TBD09: 0.94<br>[0.78; 1.13] | - |
| **Pa vs TBD09** | S Pa Q vs S Q TBD09 | 2.98 | NC | NC | S: 0.72***<br>[0.62; 0.84] | Q: 1.03<br>[0.83; 1.29] | - |
| **Sut vs 286** | S Pa Sut vs S Pa 286 | 0.96 | 1.74 | -0.79***<br>[-0.99; -0.59] | S: 1.39***<br>[1.23; 1.57] | Pa: 1.72***<br>[1.51; 1.96] | - |
| **Sut vs Q** | S Pa Sut vs S Pa Q | 0.96 | 2.98 | -2.02***<br>[-2.31; -1.74] | S: 1.12<br>[0.98; 1.28] | Pa: 1.77***<br>[1.63; 1.92] | - |
| **Sut vs TBD09** | S Pa Sut vs S Pa TBD09 | 0.96 | 2 | -1.05***<br>[-1.27; -0.82] | S: 0.89*<br>[0.81; 0.97] | Pa: 1.25***<br>[1.15; 1.36] | - |
| **Sut vs TBD11** | S Pa Sut vs S Pa TBD11 | 0.96 | 2.3 | -1.34***<br>[-1.58; -1.1] | S: 1.06<br>[0.98; 1.14] | Pa: 2.13***<br>[1.92; 2.36] | - |
| **TBD09 vs 286** | S Pa TBD09 vs S Pa 286 | 2 | 1.74 | 0.26*<br>[0; 0.51] | S: 1.56***<br>[1.38; 1.77] | Pa: 1.38***<br>[1.2; 1.57] | - |
| **TBD09 vs Q** | S Pa TBD09 vs S Pa Q | 2 | 2.98 | -0.97***<br>[-1.3; -0.65] | S: 1.27**<br>[1.1; 1.45] | Pa: 1.42***<br>[1.3; 1.55] | - |
| **TBD09 vs TBD11** | S Pa TBD09 vs S Pa TBD11 | 2 | 2.3 | -0.29*<br>[-0.58; -0.01] | S: 1.19 ***<br>[1.1; 1.3] | Pa: 1.7***<br>[1.53; 1.9] | - |
| **TBD11 vs 286** | S Pa TBD11 vs S Pa 286 | 2.3 | 1.74 | 0.55***<br>[0.28; 0.82] | S: 1.31**<br>[1.16; 1.47] | Pa: 0.81**<br>[0.7; 0.93] | - |
| **TBD11 vs Q** | S Pa TBD11 vs S Pa Q | 2.3 | 2.98 | -0.68***<br>[-1.02; -0.34] | S: 1.06<br>[0.93; 1.21] | Pa: 0.83**<br>[0.75; 0.92] | - |
| **286** | S Pa 286 Sut vs S Pa Sut | 0.82 | 0.96 | -0.13<br>[-0.31; 0.04] | S: 0.71***<br>[0.63; 0.8] | Pa: 0.78***<br>[0.72; 0.84] | Sut: 0.86*<br>[0.76; 0.96] |
|  | S Pa Q 286 vs S Pa Q | 2.47 | 2.98 | -0.51**<br>[-0.86; -0.16] | S: 1.01<br>[0.82; 1.24] | Pa: 1.12<br>[1; 1.25] | Q: 1.18<br>[0.95; 1.46] |
|  | S Q 286 TBD09 vs S Q TBD09 | 2.48 | NC | NC | S: 0.89<br>[0.71; 1.12] | Q: 0.96<br>[0.79; 1.16] | TBD09: 1.05<br>[0.87; 1.27] |
| **656** | S Pa 656 Sut vs S Pa Sut | 0.97 | 0.96 | 0.01<br>[-0.13; 0.15] | S: 1.04<br>[0.93; 1.16] | Pa: 0.75**<br>[0.63; 0.9] | Sut: 2.38***<br>[2.11; 2.7] |
| **Pa** | S Pa Q TBD09 vs S Q TBD09 | 2.12 | NC | NC | S: 0.93<br>[0.81; 1.06] | Q: 1.24*<br>[1.03; 1.48] | TBD09: 0.82*<br>[0.69; 0.98] |
| **Q** | S Pa Q 286 vs S Pa 286 | 2.47 | 1.74 | 0.72***<br>[0.44; 1.01] | S: 1.25*<br>[1.02; 1.53] | Pa: 1.08<br>[0.94; 1.26] | 286: 0.77<br>[0.56; 1.07] |
|  | S Pa Q TBD11 vs S Pa TBD11 | 1.84 | 2.3 | -0.46**<br>[-0.73; -0.19] | S: 0.97<br>[0.87; 1.09] | Pa: 0.96<br>[0.84; 1.11] | TBD11: 1.75***<br>[1.69; 1.81] |
|  | S Pa Q TBD09 vs S Pa TBD09 | 2.12 | 2 | 0.12<br>[-0.15; 0.39] | S: 1.02<br>[0.91; 1.13] | Pa: 0.9<br>[0.78; 1.04] | TBD09: 0.87<br>[0.74; 1.04] |
| **Sut** | S Pa 286 Sut vs<br>S Pa 286 | 0.82 | 1.74 | -0.92***<br>[-1.14; -0.71] | S: 0.98<br>[0.85; 1.13] | Pa: 1.34**<br>[1.17; 1.52] | 286: 1.08<br>[0.91; 1.27] |
| **TBD09** | S Pa Q TBD09 vs<br>S Pa Q | 2.12 | 2.98 | -0.86***<br>[-1.18; -0.53] | S: 1.29**<br>[1.11; 1.49] | Pa: 1.27**<br>[1.1; 1.47] | Q: 1.2<br>[0.96; 1.5] |
| **TBD11** | S Pa Q TBD11 vs<br>S Pa Q | 1.84 | 2.98 | -1.14***<br>[-1.46; -0.83] | S: 1.03<br>[0.89; 1.2] | Pa: 0.8**<br>[0.7; 0.91] | Q: 0.99<br>[0.76; 1.29] |
|  | S Q TBD11 TBD09 vs<br>S Q TBD09 | 2.11 | NC | NC | S: 0.86<br>[0.73; 1.01] | Q: 0.68**<br>[0.57; 0.83] | TBD09: 1.06<br>[0.8; 1.4] |
| **286 vs 656** | S Pa 286 Sut vs<br>S Pa 656 Sut | 0.82 | 0.97 | -0.14<br>[-0.3; 0.01] | S: 0.68***<br>[0.6; 0.78] | Pa: 1.03<br>[0.87; 1.23] | Sut: 0.36***<br>[0.32; 0.4] |
| **656 vs 830** | B Pa 656 Sut vs<br>B Pa 830 Sut | 2.52 | 2.33 | 0.19<br>[-0.07; 0.45] | B: 1.45**<br>[1.25; 1.69] | Pa: 0.9<br>[0.7; 1.15] | Sut: 1.05<br>[0.93; 1.18] |
| **Pa vs 286** | S Pa Q TBD09 vs S<br>Q 286 TBD09 | 2.12 | 2.48 | -0.36*<br>[-0.66; -0.06] | S: 1.04<br>[0.83; 1.3] | Q: 1.29*<br>[1.07; 1.56] | TBD09: 0.78*<br>[0.65; 0.94] |
| **Pa vs Del** | S Pa 656 Sut vs<br>S Del 656 Sut | 0.97 | 2.44 | -1.48***<br>[-1.63; -1.32] | S: 0.59***<br>[0.53; 0.66] | 656: 0.7*<br>[0.54; 0.9] | Sut: 0.99<br>[0.88; 1.12] |
| **Pa vs TBD09** | S Pa Q TBD11 vs<br>S Q TBD11 TBD09 | 1.84 | 2.11 | -0.27*<br>[-0.53; -0.01] | S: 0.87<br>[0.74; 1.02] | Q: 1.5**<br>[1.18; 1.91] | TBD11:<br>0.75***<br>[0.72; 0.78] |
|  | S Pa Q 286 vs<br>S Q 286 TBD09 | 2.47 | 2.48 | -0.01<br>[-0.34; 0.31] | S: 0.81<br>[0.63; 1.06] | Q: 1.27*<br>[1.06; 1.52] | 286: 0.93<br>[0.67; 1.28] |
| **Pa vs TBD11** | S Pa Q TBD09 vs<br>S Q TBD11 TBD09 | 2.12 | 2.11 | 0.01<br>[-0.26; 0.29] | S: 1.08<br>[0.93; 1.26] | Q: 1.81***<br>[1.49; 2.19] | TBD09: 0.78<br>[0.59; 1.02] |
| **Sut vs Q** | S Pa 286 Sut vs<br>S Pa Q 286 | 0.82 | 2.47 | -1.64***<br>[-1.91; -1.38] | S: 0.79*<br>[0.64; 0.97] | Pa: 1.23**<br>[1.1; 1.38] | 286: 1.39*<br>[1.01; 1.92] |
| **S vs B** | S Pa 656 Sut vs<br>B Pa 656 Sut | 0.97 | 2.52 | -1.56***<br>[-1.72; -1.39] | Pa: 2.45***<br>[1.9; 3.14] | 656: 0.69**<br>[0.56; 0.84] | Sut: 0.93<br>[0.84; 1.03] |
| **TBD09 vs 286** | S Pa Q TBD09 vs<br>S Pa Q 286 | 2.12 | 2.47 | -0.35*<br>[-0.64; -0.05] | S: 1.28*<br>[1.04; 1.56] | Pa: 1.14<br>[0.97; 1.33] | Q: 1.02<br>[0.86; 1.2] |
| **TBD11 vs 286** | S Pa Q TBD11 vs<br>S Pa Q 286 | 1.84 | 2.47 | -0.63***<br>[-0.92; -0.34] | S: 1.03<br>[0.84; 1.26] | Pa: 0.72***<br>[0.62; 0.83] | Q: 0.84<br>[0.67; 1.06] |
|  | S Q TBD11 TBD09 vs<br>S Q 286 TBD09 | 2.11 | 2.48 | -0.37*<br>[-0.68; -0.07] | S: 0.96<br>[0.76; 1.22] | Q: 0.71**<br>[0.59; 0.87] | TBD09: 1.01<br>[0.76; 1.34] |
| **TBD11 vs**<br>**TBD09** | S Pa Q TBD11 vs<br>S Pa Q TBD09 | 1.84 | 2.12 | -0.28*<br>[-0.54; -0.03] | S: 0.8**<br>[0.71; 0.91] | Pa: 0.63***<br>[0.53; 0.74] | Q: 0.83<br>[0.65; 1.05] |

### Assessment of model prediction performance through a simulation-based approach

To obtain prediction intervals for the combinations included in the two RMM studies of interest for this paper, 200 relapse profiles were simulated by mimicking the covariate distribution of the studies included into the model (i.e. inoculum) and assuming the estimated inter-study variability (η_1_=0.045). Figure S1 shows median and 90% prediction intervals for each combination. Overall, most of the observed data are included in the 90% prediction intervals, indicating good prediction performance of the model especially for comparator BPaMZ for which data coming from different studies are available.

**Figure S1-.**
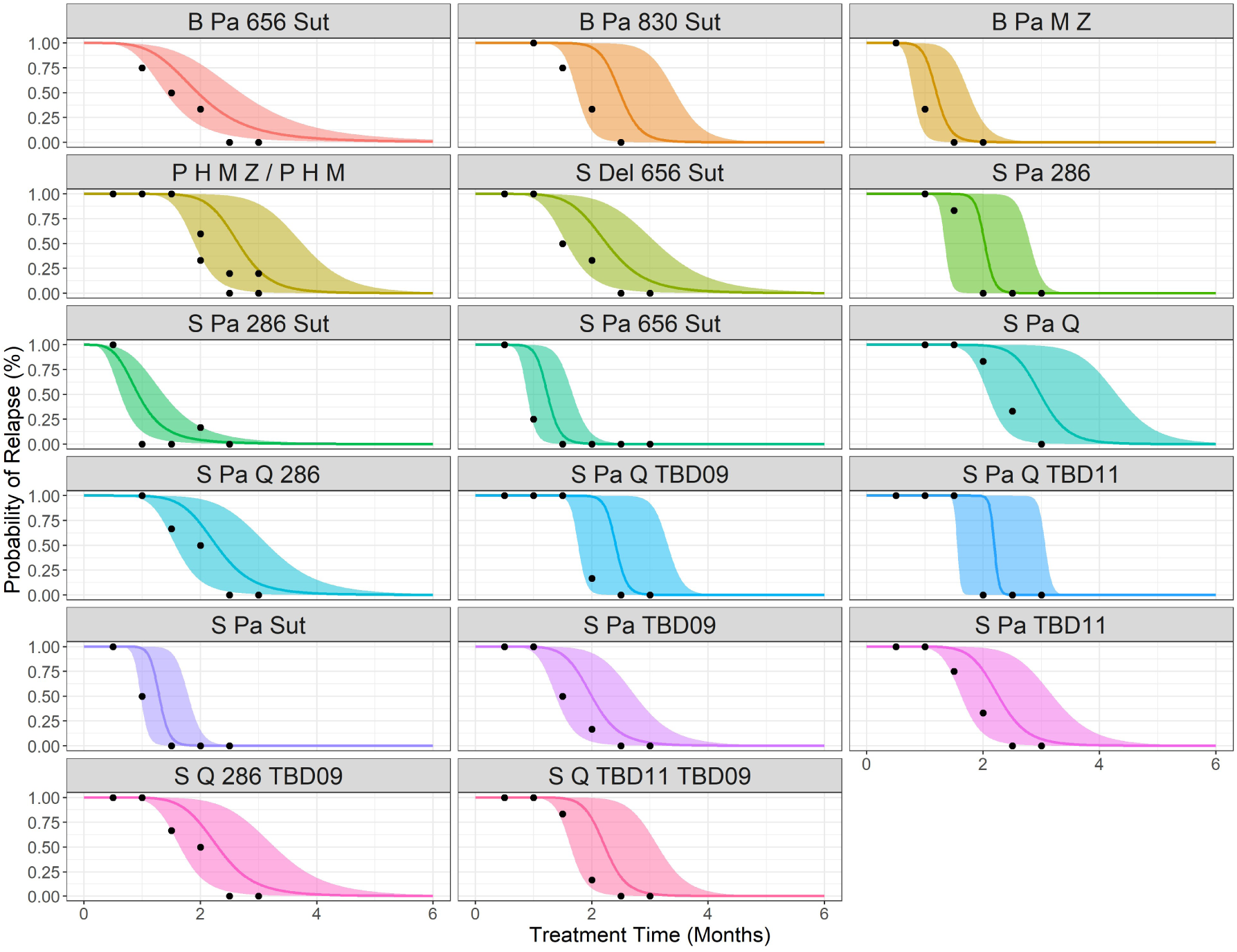
Prediction intervals for Study 1 and Study 2 combinations obtained with model-based simulation. For each combination, median (solid lines) and 90% confidence interval (shaded areas) are reported, together with relapse data observed in the two studies (black dots).

**Figure S2.**
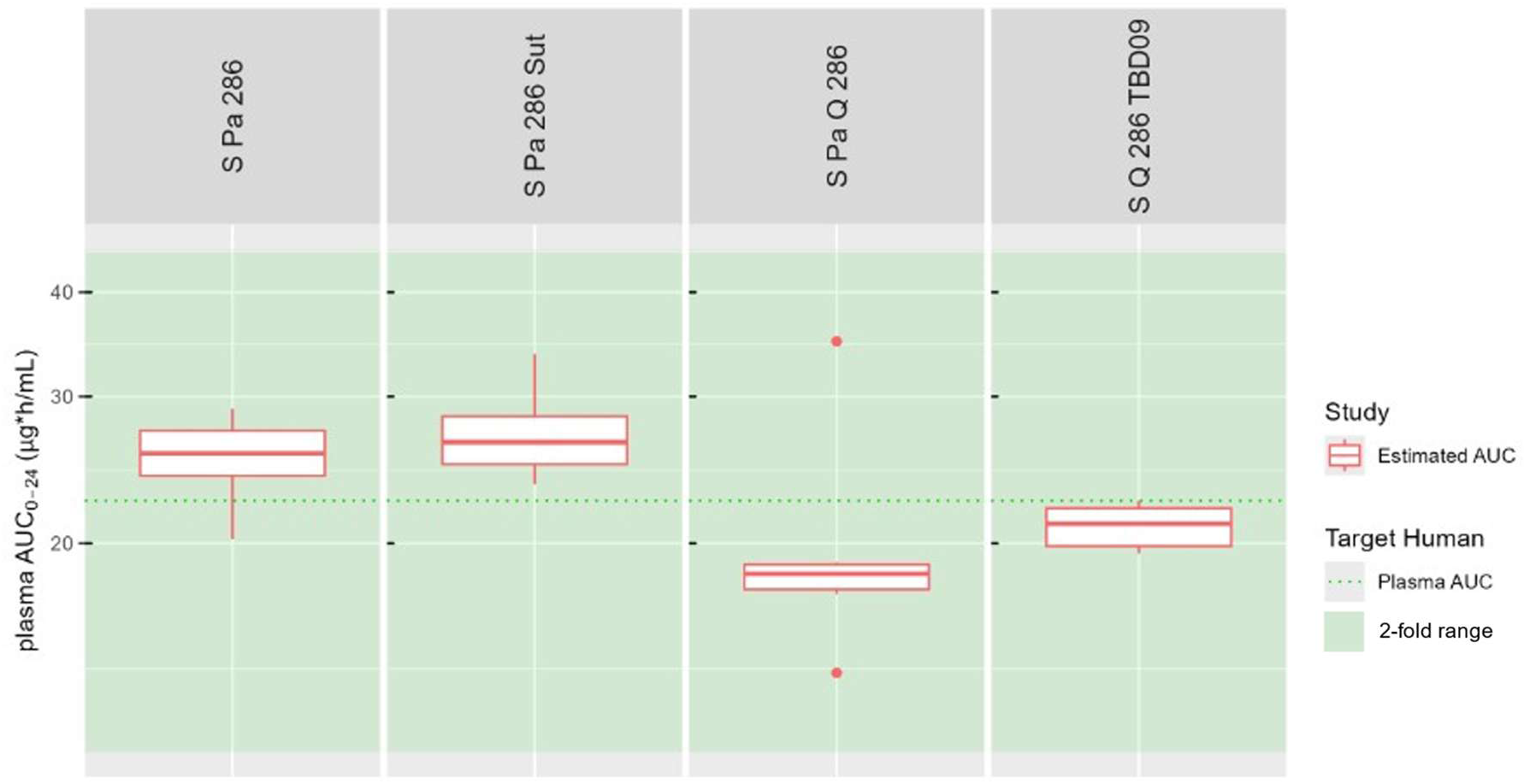
Distribution of plasma AUC_0-24_ of 286 from popPK analysis for combinations sharing 286.

**Figure S3.**
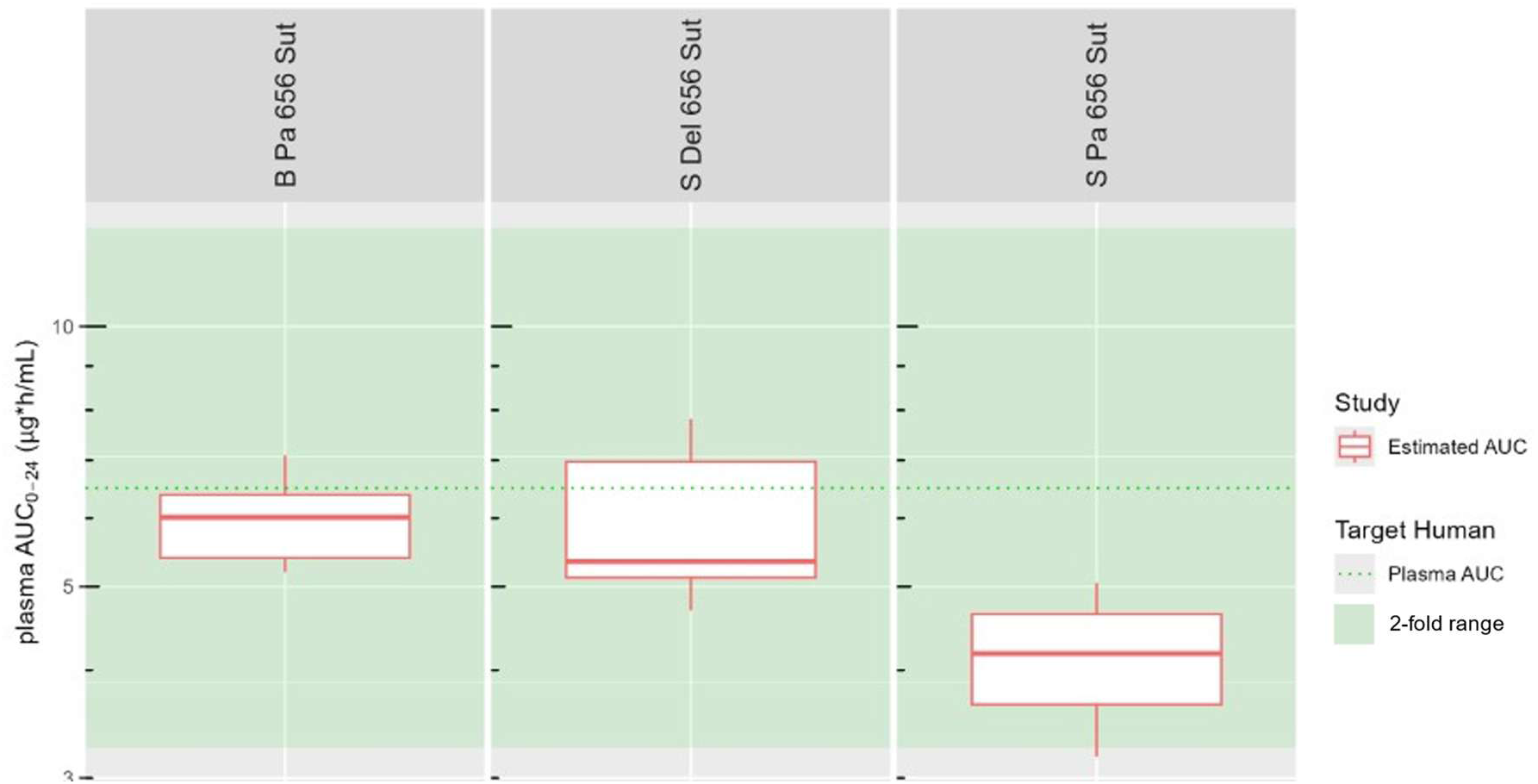
Distribution of plasma AUC_0-24_ of 656 from popPK analysis for combinations sharing 656.

**Figure S4.**
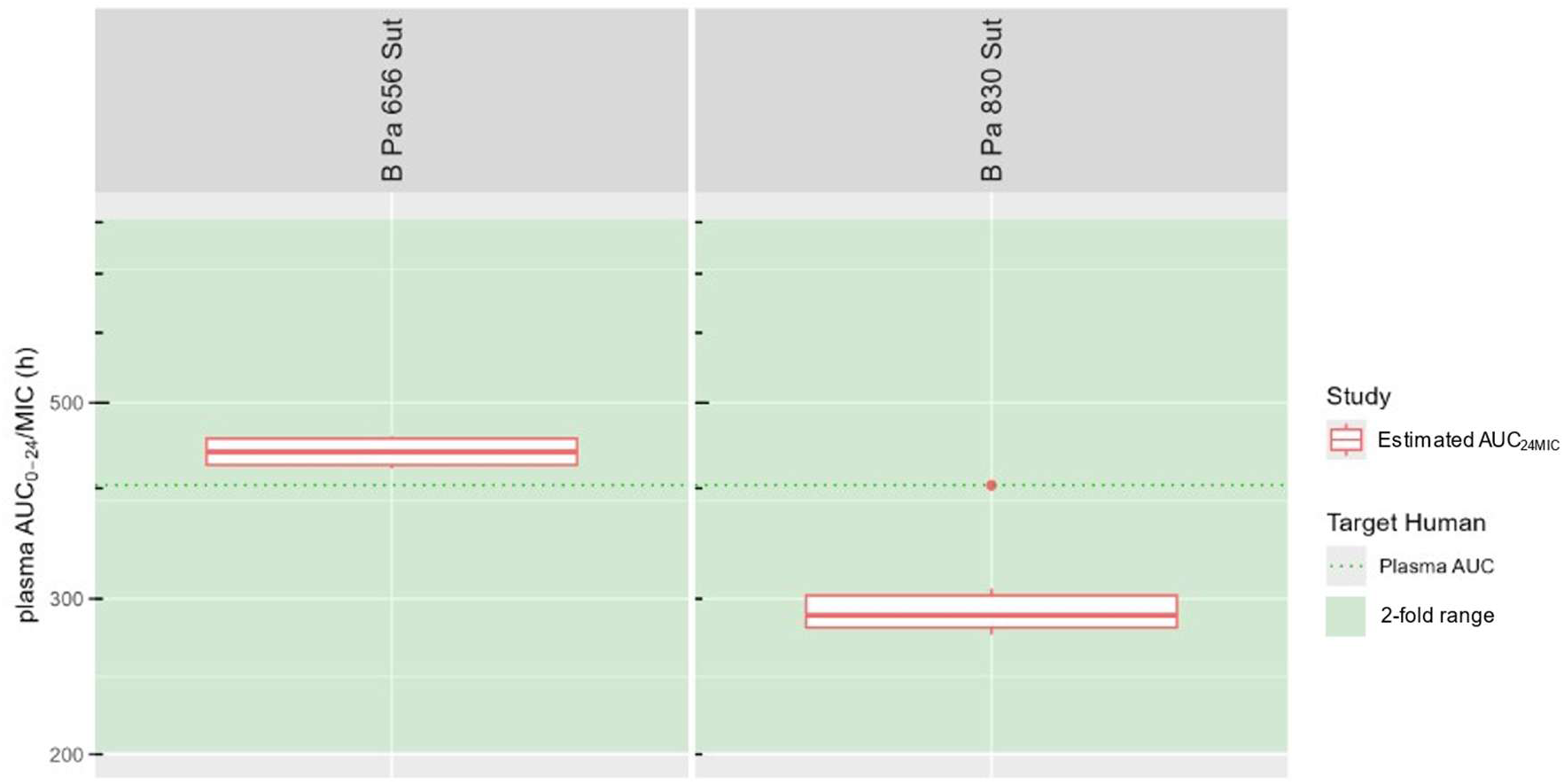
Distribution of plasma AUC_24MIC_ of B from popPK analysis for combinations sharing B.

**Figure S5.**
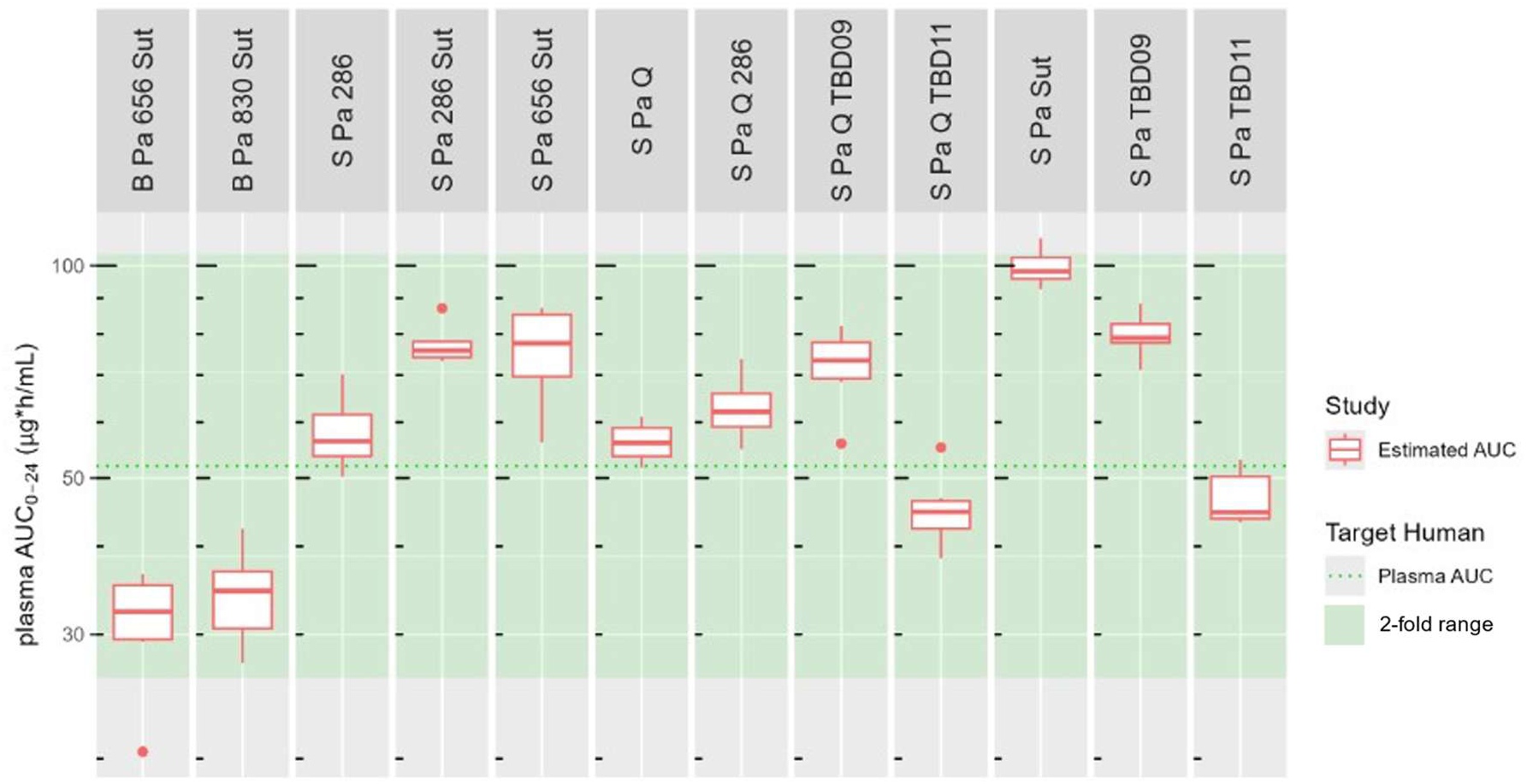
Distribution of plasma AUC_0-24_ of Pa from popPK analysis for combinations sharing Pa.

**Figure S6.**
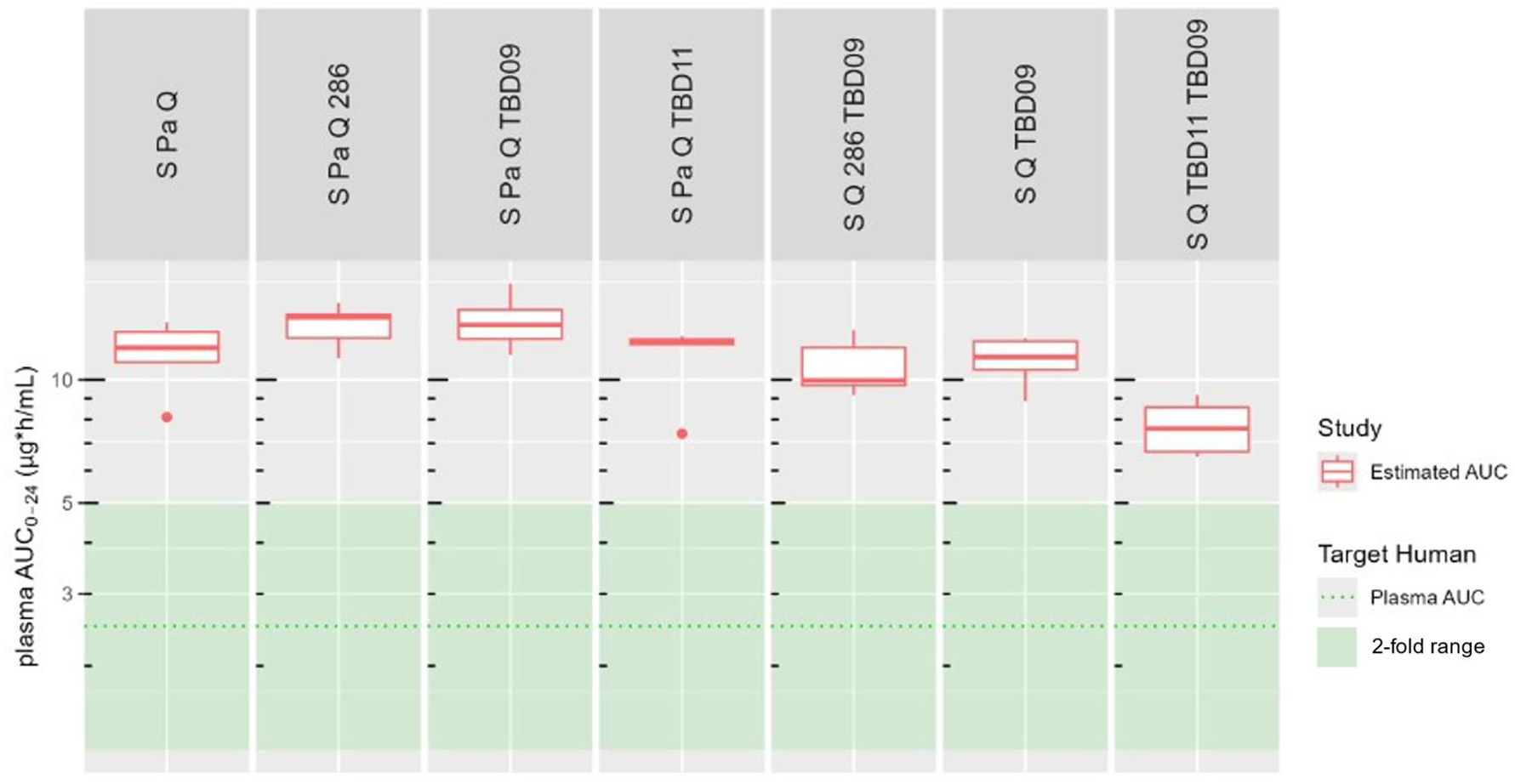
Distribution of plasma AUC_0-24_ of Q from popPK analysis for combinations sharing Q.

**Figure S7.**
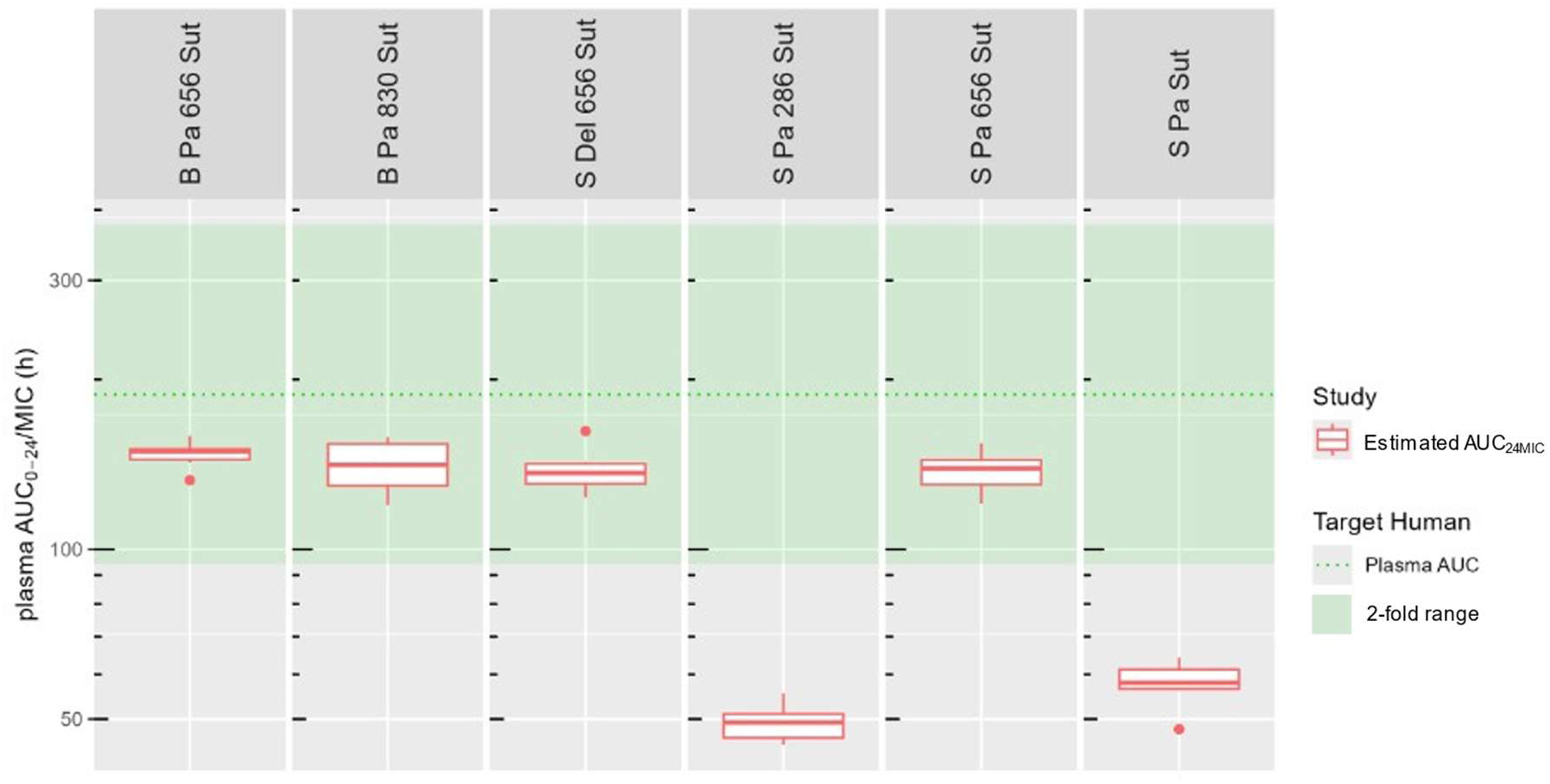
Distribution of plasma AUC_24MIC_ of Sut from popPK analysis for combinations sharing Sut.

**Figure S8.**
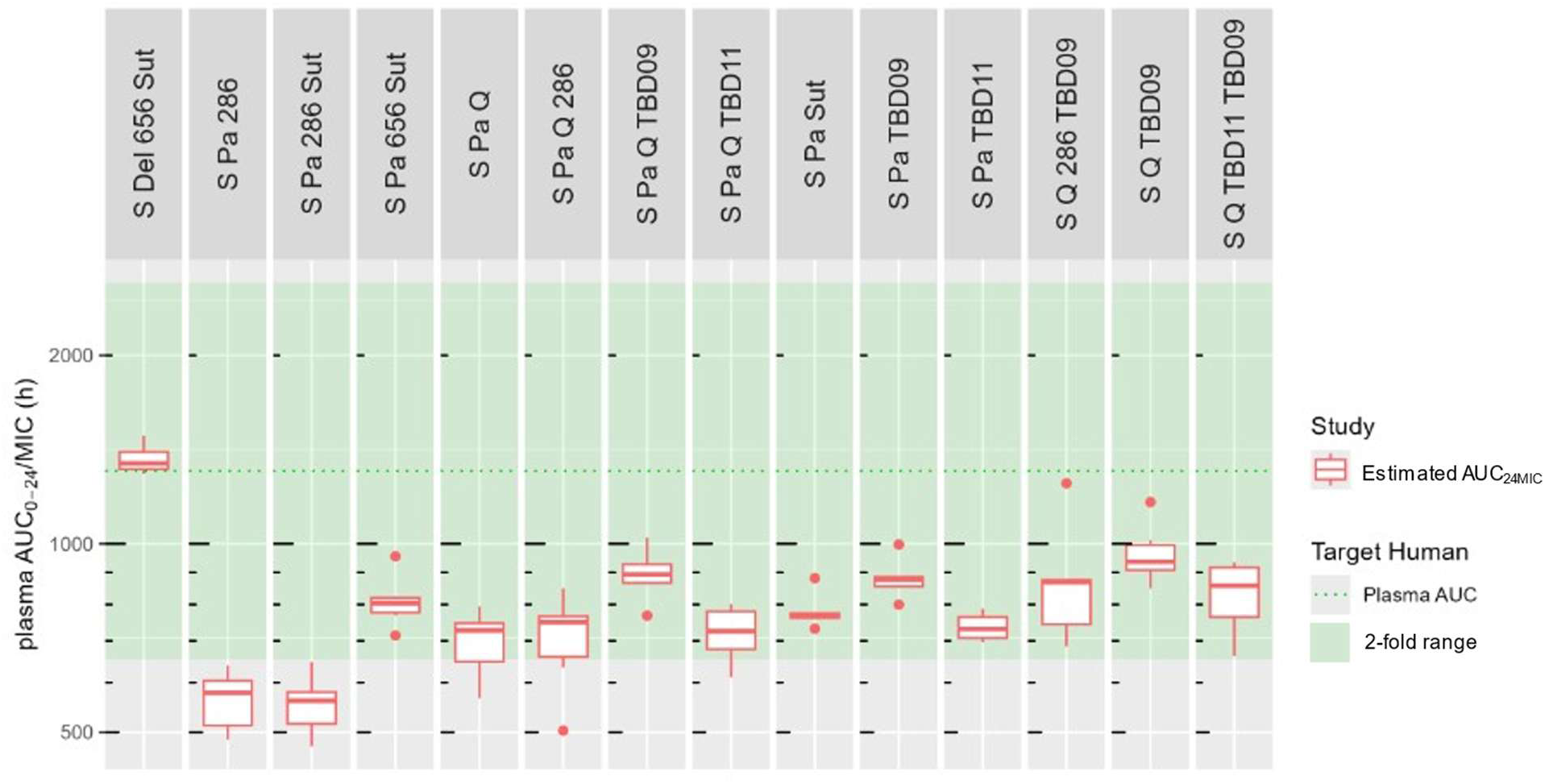
Distribution of plasma AUC_24MIC_ of S from popPK analysis for combinations sharing S.

**Figure S9.**
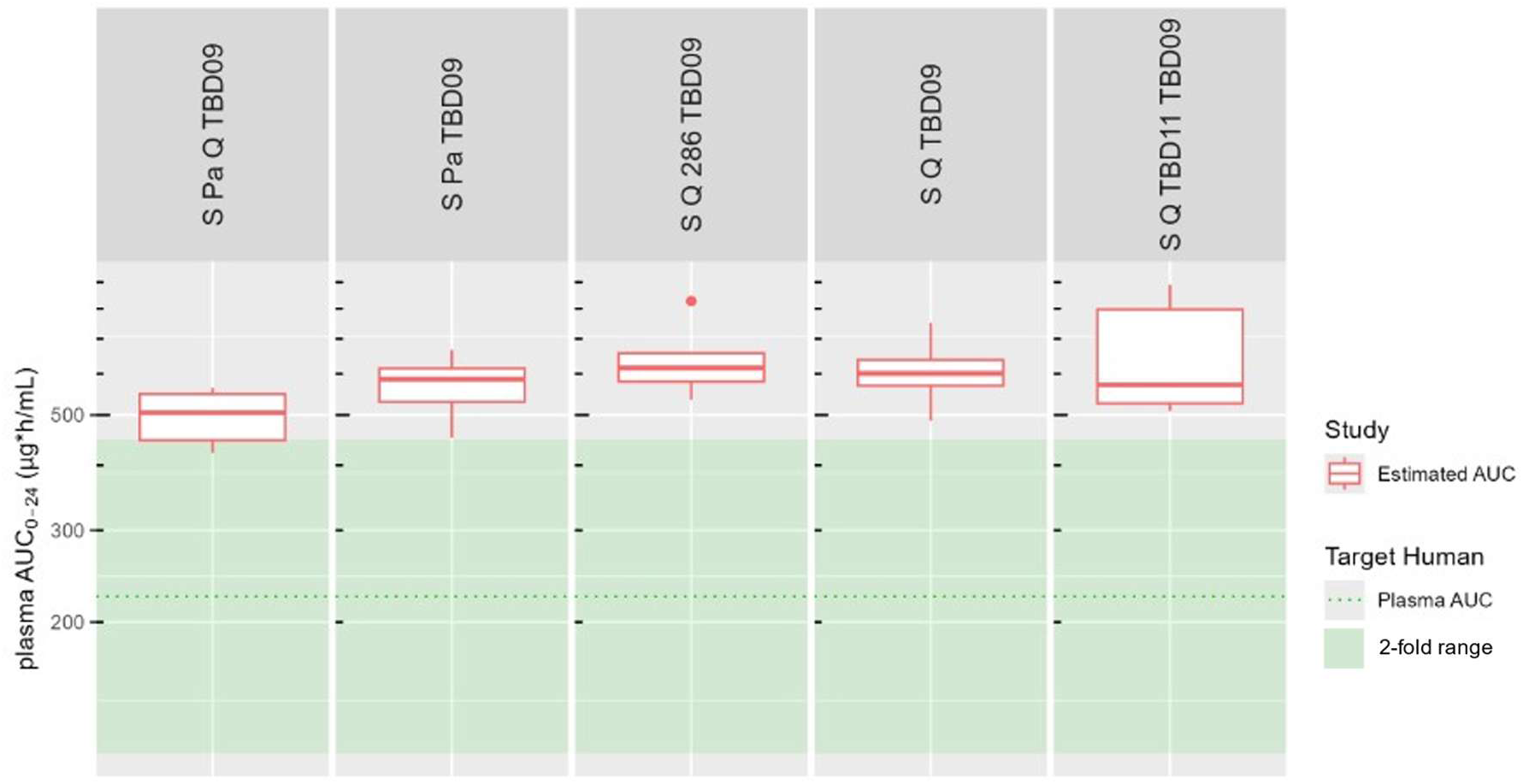
Distribution of plasma AUC_0-24_ of TBD09 from popPK analysis for combinations sharing TBD09.

**Figure S10.**
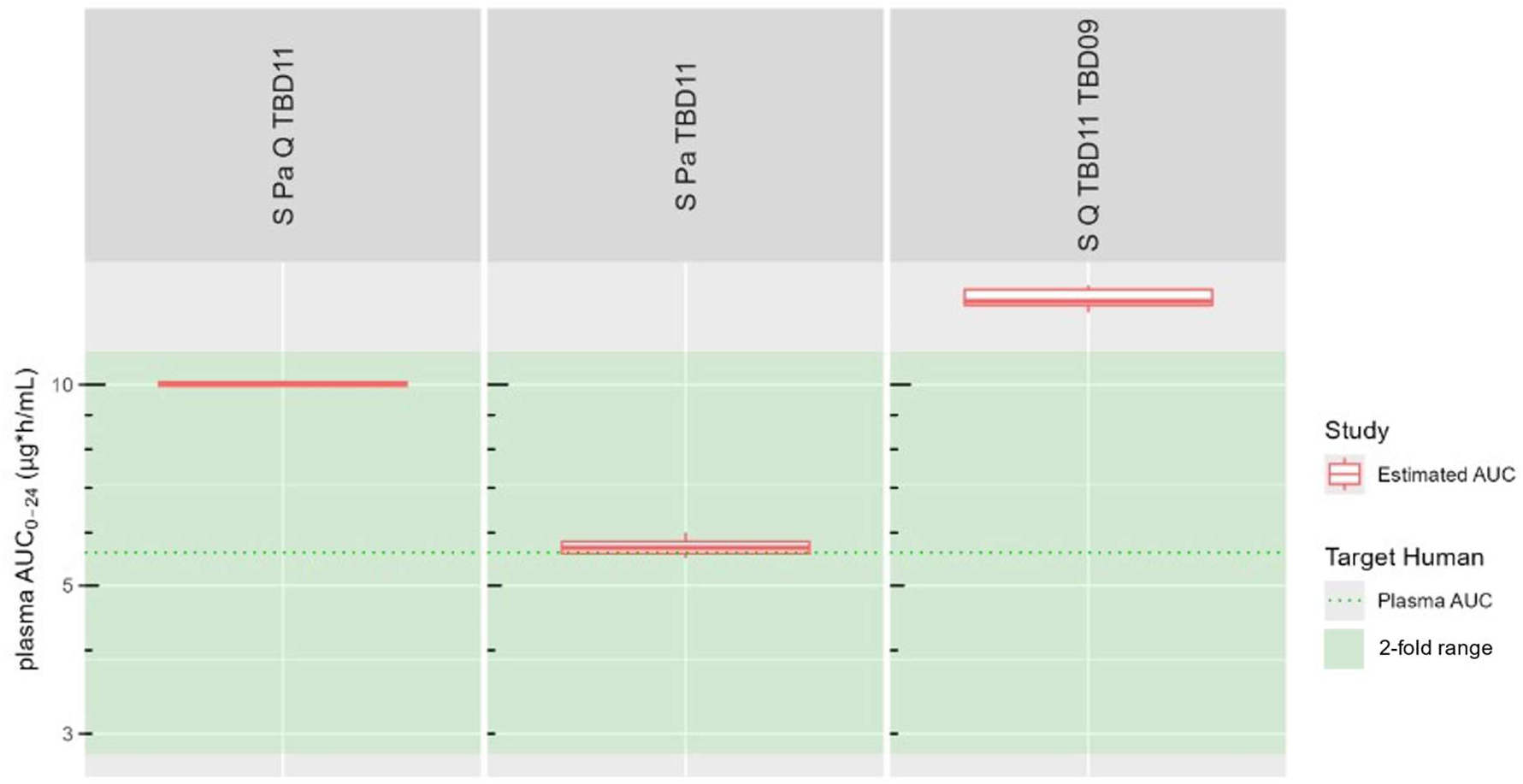
Distribution of plasma AUC_0-24_ of TBD11 from popPK analysis for combinations sharing TBD11.

## REFERENCES

[1] World Health Organization, Global tuberculosis report 2024

[2] Cox H, Dickson-Hall L, Ndjeka N, Van’t Hoog A, Grant A, Cobelens F, Stevens W, Nicol M. 2017. Delays and loss to follow-up before treatment of drug-resistant tuberculosis following implementation of Xpert MTB/RIF in South Africa: a retrospective cohort study. PLoS Med. 14(2):e1002238. doi: 10.1371/journal.pmed.1002238.

[3] Lienhardt C, Dooley KE, Nahid P, Wells C, Ryckman TS, Kendall EA, Davies G, Brigden G, Churchyard G, Cirillo DM, Di Meco E, Gopinath R, Mitnick C, Scott C, Amanullah F, Bansbach C, Boeree M, Campbell M, Conradie F, Crook A, Daley CL, Dheda K, Diacon A, Gebhard A, Hanna D, Heinrich N, Hesseling A, Holtzman D, Jachym M, Kim P, Lange C, McKenna L, Meintjes G, Ndjeka N, Nhung NV, Nyang’wa BT, Paton NI, Rao R, Rich M, Savic R, Schoeman I, Makokotlela BS, Spigelman M, Sun E, Svensson E, Tisile P, Varaine F, Vernon A, Diul MY, Kasaeva T, Zignol M, Gegia M, Mirzayev F, Schumacher SG. 2024. Target regimen profiles for tuberculosis treatment. Bull World Health Organ. 102(8):600–607. doi: 10.2471/BLT.24.291881. Epub 2024 May 28.

[4] WHO Target regimen profiles for tuberculosis treatment. Nov 2, 2023. https://www.who.int/publications-detail-redirect/9789240081512 [DOI]

[5] Hards K, Robson JR, Berney M, Shaw L, Bald D, Koul A, Andries K, Cook GM. 2015. Bactericidal mode of action of bedaquiline. J Antimicrob Chemother. 70(7):2028–37. doi: 10.1093/jac/dkv054. Epub 2015 Mar 8. PMID: 25754998

[6] Conradie F, Bagdasaryan TR, Borisov S, Howell P, Mikiashvili L, Ngubane N, Samoilova A, Skornykova S, Tudor E, Variava E, Yablonskiy P, Everitt D, Wills GH, Sun E, Olugbosi M, Egizi E, Li M, Holsta A, Timm J, Bateson A, Crook AM, Fabiane SM, Hunt R, McHugh TD, Tweed CD, Foraida S, Mendel CM, Spigelman M; ZeNix Trial Team. 2022. Bedaquiline-Pretomanid-Linezolid Regimens for Drug-Resistant Tuberculosis. N Engl J Med 387(9): 810–823. 10.1056/NEJMoa2119430

[7] Khoshnood S, Taki E, Sadeghifard N, Kaviar VH, Haddadi MH, Farshadzadeh Z, Kouhsari E, Goudarzi M, Heidary M. 2021. Mechanism of Action, Resistance, Synergism, and Clinical Implications of Delamanid Against Multidrug-Resistant *Mycobacterium tuberculosis*. Front Microbiol 12: 717045. 10.3389/fmicb.2021.717045

[8] Hariguchi N, Chen X, Hayashi Y, Kawano Y, Fujiwara M, Matsuba M, Shimizu H, Ohba Y, Nakamura I, Kitamoto R, Shinohara T, Uematsu Y, Ishikawa S, Itotani M, Haraguchi Y, Takemura I, Matsumoto M. 2020. OPC-167832, a Novel Carbostyril Derivative with Potent Antituberculosis Activity as a DprE1 Inhibitor. Antimicrob Agents Chemother 64(6). 10.1128/aac.02020-19

[9] Tenero D, Derimanov G, Carlton A, Tonkyn J, Davies M, Cozens S, Gresham S, Gaudion A, Puri A, Muliaditan M, Rullas-Trincado J, Mendoza-Losana A, Skingsley A, Barros-Aguirre D. 2019. First-Time-in-Human Study and Prediction of Early Bactericidal Activity for GSK3036656, a Potent Leucyl-tRNA Synthetase Inhibitor for Tuberculosis Treatment. Antimicrob Agents Chemother 63(8). 10.1128/aac.00240-19

[10] Brown KL, Wilburn KM, Montague CR, Grigg JC, Sanz O, Pérez-Herrán E, Barros D, Ballell L, VanderVen BC, Eltis LD. 2023. Cyclic AMP-Mediated Inhibition of Cholesterol Catabolism in Mycobacterium tuberculosis by the Novel Drug Candidate GSK2556286. Antimicrob Agents Chemother 67(1): e0129422. 10.1128/aac.01294-22

[11] Bruinenberg P, Nedelman J, Yang TJ, Pappas F, Everitt D. 2022. Single Ascending-Dose Study To Evaluate the Safety, Tolerability, and Pharmacokinetics of Sutezolid in Healthy Adult Subjects. Antimicrob Agents Chemother 66(4): e0210821. 10.1128/aac.02108-21

[12] Sordello S, Tagliavini A, Boulenc X, Brock L, Zannoni S, Roversi C, Visentin R, Metcalf D, Federico D, Modolo S, Calusi G, Petterlini R, Golovkine G, Pascal C, Huc Claustre E, Vahlas Z, Pergher M, Mdluli K, Levi M, Black T, Bates R, Liu Y, Aguilar Perez C, Hermann D, Flood D, Upton A.M. 2026 Nonclinical pharmacokinetics and relative efficacy of the first 25 novel tuberculosis drug combinations from the PAN-TB consortium: Use of the BALB/c relapsing mouse model and combination pharmacokinetics within a modeling-based framework. Submitted AAC

[13] Lombardi A, Pappas F, Nedelman J, Hickman D, Jaw-Tsai S, Olugbosi M, Bruinenberg P, Beumont M, Sun E. 2024 Pharmacokinetics and safety of TBAJ-876, a novel antimycobacterial diarylquinoline, in healthy subjects. Antimicrob Agents Chemother. 2024 Oct 8;68(10):e0061324. doi: 10.1128/aac.00613-24. Epub 2024 Aug 28. PMID: 39194204

[14] Crowley BM, Boshoff HI, Boving A, Tan VY, Zhu J, Hoyt F, Miller RR, Ehrhart J, Boyce CW, Young K, Nantermet PG, Su J, Yang L, Painter RE, Corcoran EB, Hoar JL, Oh S, Holtzman DL, Levi M, Anderson A, Otieno MA, Zimmerman M, Kaya F, Massoudi LM, Ramey ME, Bauman AA, Lenaerts AJ, Roberston GT, Dartois V, Wells CD, Barry CE 3rd, Olsen DB. 2026 Discovery and development of a new oxazolidinone with reduced toxicity for the treatment of tuberculosis. Nat Med. 2026 Jan 13. doi: 10.1038/s41591-025-04164-x. Online ahead of print.

[15] Wilburn KM, Montague CR, Qin B, Woods AK, Love MS, McNamara CW, Schultz PG, Southard TL, Huang L, Petrassi HM, VanderVen BC. 2022. Pharmacological and genetic activation of cAMP synthesis disrupts cholesterol utilization in Mycobacterium tuberculosis. PLoS Pathog. 2022 Feb 8;18(2):e1009862. doi: 10.1371/journal.ppat.1009862. eCollection 2022 Feb. PMID: 35134095

[16] Martinez G, Tolentino K, Sukheja P, Webb J, McNamara CW, Chatterjee AK, Yang B. 2025. Novel isoxazole thiophene-containing compounds active against Mycobacterium tuberculosis. Bioorg Med Chem Lett. 2025 Apr 15;119:130108. doi: 10.1016/j.bmcl.2025.130108. Epub 2025 Jan 23. PMID: 39863083

[17] Li SY, Converse PJ, Betoudji F, Lee J, Mdluli K, Upton A, Fotouhi N, Nuermberger EL. 2023. Next-Generation Diarylquinolines Improve Sterilizing Activity of Regimens with Pretomanid and the Novel Oxazolidinone TBI-223 in a Mouse Tuberculosis Model. Antimicrob Agents Chemother. 2023 Apr 18;67(4):e0003523. doi: 10.1128/aac.00035-23. Epub 2023 Mar 15

[18] Dorman SE, Nahid P, Kurbatova EV, Phillips PPJ, Bryant K, Dooley KE, Engle M, Goldberg SV, Phan HTT, Hakim J, Johnson JL, Lourens M, Martinson NA, Muzanyi G, Narunsky K, Nerette S, Nguyen NV, Pham TH, Pierre S, Purfield AE, Samaneka W, Savic RM, Sanne I, Scott NA, Shenje J, Sizemore E, Vernon A, Waja Z, Weiner M, Swindells S, Chaisson RE, Tuberculosis Trials Consortium. 2021. Four-month rifapentine regimens with or without moxifloxacin for tuberculosis. N Engl J Med 384:1705–1718. 10.1056/NEJMoa2033400.

[19] Cevik M, Thompson LC, Upton C, Rolla VC, Malahleha M, Mmbaga B, Ngubane N, Abu Bakar Z, Rassool M, Variava E, Dawson R, Staples S, Lalloo U, Louw C, Conradie F, Eristavi M, Samoilova A, Skornyakov SN, Ntinginya NE, Haraka F, Praygod G, Mayanja-Kizza H, Caoili J, Balanag V, Dalcolmo MP, McHugh T, Hunt R, Solanki P, Bateson A, Crook AM, Fabiane S, Timm J, Sun E, Spigelman M, Sloan DJ, Gillespie SH; SimpliciTB Consortium. 2024. Bedaquiline-pretomanid-moxifloxacin-pyrazinamide for drug-sensitive and drug-resistant pulmonary tuberculosis treatment: a phase 2c, open-label, multicentre, partially randomised controlled trial. The Lancet Infectious Diseases 24(9): 1003–1014. 10.1016/S1473-3099(24)00223-8

[20] Heinrich N, Manyama C, Koele SE, Mpagama S, Mhimbira F, Sebe M, Wallis RS, Ntinginya N, Liyoyo A, Huglin B, Minja LT, Wagnerberger L, Stoycheva K, Zumba T, Noreña I, Peter DD, Makkan H, Sloan DJ, Brake LT, Schildkraut J, Aarnoutse RE, McHugh TD, Wildner L, Boeree M, Aldana BH, Phillips PPJ, Hoelscher M, Svensson EM; PanACEA consortium. 2025. Sutezolid in combination with bedaquiline, delamanid, and moxifloxacin for pulmonary tuberculosis (PanACEA-SUDOCU-01): a prospective, open-label, randomised, phase 2b dose-finding trial. Lancet Infect Dis. 2025 Nov;25(11):1208–1218. doi: 10.1016/S1473-3099(25)00213-0. Epub 2025 Jul 8.

[21] Tenero D, Derimanov G, Carlton A, Tonkyn J, Davies M, Cozens S, Gresham S, Gaudion A, Puri A, Muliaditan M, Rullas-Trincado J, Mendoza-Losana A, Skingsley A, Barros-Aguirre D. 2019. First-Time-in-Human Study and Prediction of Early Bactericidal Activity for GSK3036656, a Potent Leucyl-tRNA Synthetase Inhibitor for Tuberculosis Treatment. Antimicrob Agents Chemother. 2019 Jul 25;63(8):e00240–19. doi:10.1128/AAC.00240-19. Print 2019 Aug.

[22] Diacon AH, Barry CE 3rd, Carlton A, Chen RY, Davies M, de Jager V, Fletcher K, Koh GCKW, Kontsevaya I, Heyckendorf J, Lange C, Reimann M, Penman SL, Scott R, Maher-Edwards G, Tiberi S, Vlasakakis G, Upton CM, Aguirre DB. 2024 A first-in-class leucyl-tRNA synthetase inhibitor, ganfeborole, for rifampicin-susceptible tuberculosis: a phase 2a open-label, randomized trial. Nat Med. 2024 Mar;30(3):896–904. doi: 10.1038/s41591-024-02829-7. Epub 2024 Feb 16.PMID: 38365949

[23] Nuermberger EL, Martínez-Martínez MS, Sanz O, Urones B, Esquivias J, Soni H, Tasneen R, Tyagi S, Li SY, Converse PJ, Boshoff HI, Robertson GT, Besra GS, Abrahams KA, Upton AM, Mdluli K, Boyle GW, Turner S, Fotouhi N, Cammack NC, Siles JM, Alonso M, Escribano J, Lelievre J, Rullas-Trincado J, Pérez-Herrán E, Bates RH, Maher-Edwards G, Barros D, Ballell L, Jiménez E. 2022. GSK2556286 Is a Novel Antitubercular Drug Candidate Effective *In Vivo* with the Potential To Shorten Tuberculosis Treatment. Antimicrob Agents Chemother. 2022 Jun 21;66(6):e0013222. doi: 10.1128/aac.00132-22. Epub 2022 May 24.

[24] Li S-Y, Tyagi S, Soni H, Betoudji F, Converse PJ, Mdluli K, Upton AM, Fotouhi N, Barros-Aguirre D, Ballell L, Jimenez-Navarro E, Nuermberger EL. 2024. Bactericidal and sterilizing activity of novel regimens combining bedaquiline or TBAJ-587 with GSK2556286 and TBA-7371 in a mouse model of tuberculosis.Antimicrob Agents Chemother. 2024 Apr 3;68(4):e0156223. doi: 10.1128/aac.01562-23. Epub 2024 Feb 20. PMID: 38376228

[25] Sordello S, Brock L, Tagliavini A, Federico D, Boulenc X, Pergher M, Huc Claustre E, Metcalf D, Walter ND, Robertson GT, Clary J, Berg A, Mdluli K, Hermann D, Flood D, Upton AM. 2026 A modeling-based framework to evaluate forgiveness of tuberculosis treatment in a BALB/c relapsing mouse model. Antimicrob Agents Chemother. 2026 Feb 4;70(2):e0110925. doi: 10.1128/aac.01109-25. Epub 2026 Jan 15. PMID: 41537590

[26] Li, W., Vazvaei-Smith, F., Dear, G., Boer, J., Cuyckens, F., Fraier, D., Liang, Y., Lu, D., Mangus, H., Moliner, P., Pedersen, M. L., Romeo, A. A., Spracklin, D. K., Wagner, D. S., Winter, S., & Xu, X. S. (2024). Metabolite Bioanalysis in Drug Development: Recommendations from the IQ Consortium Metabolite Bioanalysis Working Group. Clinical pharmacology and therapeutics, 115(5), 939–953. 10.1002/cpt.3144

[27] MacLeod, A. K., Coquelin, K. S., Huertas, L., Simeons, F. R. C., Riley, J., Casado, P., Guijarro, L., Casanueva, R., Frame, L., Pinto, E. G., Ferguson, L., Duncan, C., Mutter, N., Shishikura, Y., Henderson, C. J., Cebrian, D., Wolf, C. R., & Read, K. D. (2024). Acceleration of infectious disease drug discovery and development using a humanized model of drug metabolism. Proceedings of the National Academy of Sciences of the United States of America, 121(7), e2315069121. 10.1073/pnas.2315069121

[28] Rouan, M. C., Lounis, N., Gevers, T., Dillen, L., Gilissen, R., Raoof, A., & Andries, K. (2012). Pharmacokinetics and pharmacodynamics of TMC207 and its N-desmethyl metabolite in a murine model of tuberculosis. Antimicrobial agents and chemotherapy, 56(3), 1444–1451. 10.1128/AAC.00720-11

[29] Wallis, R. S., Dawson, R., Friedrich, S. O., Venter, A., Paige, D., Zhu, T., Silvia, A., Gobey, J., Ellery, C., Zhang, Y., Eisenach, K., Miller, P., & Diacon, A. H. (2014). Mycobactericidal activity of sutezolid (PNU-100480) in sputum (EBA) and blood (WBA) of patients with pulmonary tuberculosis. PloS one, 9(4), e94462. 10.1371/journal.pone.0094462

[30] Berg A, Clary J, Hanna D, Nuermberger E, Lenaerts A, Ammerman N, Ramey M, Hartley D, Hermann D. 2022. Model-Based Meta-Analysis of Relapsing Mouse Model Studies from the Critical Path to Tuberculosis Drug Regimens Initiative Database. Antimicrob Agents Chemother. 2022 Mar 15;66(3):e0179321. doi: 10.1128/AAC.01793-21. Epub 2022 Jan 31.

[31] Li SY, Tasneen R, Tyagi S, Soni H, Converse PJ, Mdluli K, Nuermberger EL. 2017. Bactericidal and Sterilizing Activity of a Novel Regimen with Bedaquiline, Pretomanid, Moxifloxacin, and Pyrazinamide in a Murine Model of Tuberculosis. Antimicrob Agents Chemother 61(9). 10.1128/aac.00913-17

[32] Xu J, Li SY, Almeida DV, Tasneen R, Barnes-Boyle K, Converse PJ, Upton AM, Mdluli K, Fotouhi N, Nuermberger EL. 2019. Contribution of Pretomanid to Novel Regimens Containing Bedaquiline with either Linezolid or Moxifloxacin and Pyrazinamide in Murine Models of Tuberculosis. Antimicrob Agents Chemother. 2019 Apr 25;63(5):e00021–19. doi: 10.1128/AAC.00021-19. Print 2019 May.PMID: 30833432

[33] Walter ND, Born SEM, Robertson GT, Reichlen M, Dide-Agossou C, Ektnitphong VA, Rossmassler K, Ramey ME, Bauman AA, Ozols V, Bearrows SC, Schoolnik G, Dolganov G, Garcia B, Musisi E, Worodria W, Huang L, Davis JL, Nguyen NV, Nguyen HV, Nguyen ATV, Phan H, Wilusz C, Podell BK, Sanoussi ND, de Jong BC, Merle CS, Affolabi D, McIlleron H, Garcia-Cremades M, Maidji E, Eshun-Wilson F, Aguilar-Rodriguez B, Karthikeyan D, Mdluli K, Bansbach C, Lenaerts AJ, Savic RM, Nahid P, Vásquez JJ, Voskuil MI. 2021. Mycobacterium tuberculosis precursor rRNA indicates treatment-shortening activity of drugs and regimens. Nature Communications 12(1):2899.

